# A synthetic receptor platform enables rapid and portable monitoring of liver dysfunction via engineered bacteria

**DOI:** 10.1101/2021.03.24.436753

**Authors:** Hung-Ju Chang, Ana Zuniga, Ismael Conejero, Peter L. Voyvodic, Jerome Gracy, Elena Fajardo-Ruiz, Martin Cohen-Gonsaud, Guillaume Cambray, Georges-Philippe Pageaux, Magdalena Meszaros, Lucy Meunier, Jerome Bonnet

## Abstract

Bacterial biosensors, or bactosensors, are promising field-deployable agents for medical and environmental diagnostics. However, the lack of scalable frameworks to systematically program ligand detection limits their applications. Here we present a synthetic receptor platform, termed EMeRALD (**E**ngineered **M**odularized **R**eceptors **A**ctivated via **L**igand-induced **D**imerization) which supports the modular assembly of sensing modules onto a high-performance, generic signaling scaffold controlling gene expression in *E. coli*. We applied EMeRALD to detect bile salts, a biomarker of liver dysfunction, by repurposing sensing modules from enteropathogenic *Vibrio* species. We improved the sensitivity and lowered the limit-of-detection of the sensing module by directed evolution. We then engineered a colorimetric bactosensor detecting pathological bile salt levels in serum from patients having undergone liver transplant, providing an output detectable by the naked-eye. The EMeRALD technology enables functional exploration of natural sensing modules and rapid engineering of synthetic receptors for diagnostics, environmental monitoring, and control of therapeutic microbes.

## INTRODUCTION

Early disease detection and monitoring of chronic pathologies help reduce mortality and improve patients’ quality of life^1,2^. In that context, *in vitro* diagnostics technologies play key roles at different stages of the healthcare chain^3^. However, many diagnostics technologies require heavy-equipments, are expensive, and necessitate trained personnel, limiting testing to centralized facilities such as hospitals. Yet, as exemplified by glucose monitoring for diabetes, field-deployable diagnostic devices can tremendously improve patient healthcare, follow-up, and self-reliance^4,5^. Robust, scalable, and cost-effective biosensing technologies for field-deployable diagnostics have thus been under intense research interest over the past decade^6^.

Bacteria must sense and respond to myriad chemical and physical signals to survive and reproduce, and are thus ideal candidates for engineering biosensors. Whole-cell biosensors (WCB) are genetically modified living cells that detect molecules of interest generally using a transcription factor regulated by the ligand of interest, and activating transcription of a reporter gene^7^. While explored since the dawn of genetic engineering^8^, recent advances in synthetic biology have improved whole-cell biosensor robustness, sensitivity, signal-to-noise ratio, and signal-processing capabilities, supporting their use in complex media like wastewater and clinical samples^9,10,11^. With these new capabilities, whole-cell biosensors have the potential to provide miniaturized, field-deployable, diagnostic devices capable of multiplexed detection and sophisticated computation^10,11^. As a self-manufacturing and biodegradable platform, biology provides a cost-effective and environmentally friendly alternative to traditional diagnostic methods. The self-manufacturing nature of biology also offers a unique advantage for low-resource settings and remote, highly-constrained conditions, such as those found in space exploration^12^. Despite all these advantages, the scope of application for bacterial biosensors is limited by the difficulty to generate novel sensors detecting biomarkers of interest. Although significant progress has been recently made^13^, scalable platforms to rapidly generate new receptors are needed.

Liver disease includes dozens of pathologies such as cirrhosis, hepatitis, liver cancer, hepatobiliary problems such as cholangitis, and drug-induced liver injury^14–18^. Liver disease is a global healthcare burden accounting for 2 million deaths per year, and impacts the quality of life of millions of people worldwide^19^. Liver is the second most common transplanted organ, but only 10% of needs are met. Currently, liver disease diagnostics and monitoring is performed by assessing a panel of biomarkers^20^. However, current methods for *in vitro* diagnostics are only available in hospitals and testing laboratories, limiting the frequency of monitoring for patients. In addition, most biomarkers appear at late disease stages (when significant cellular damage has already occurred) and lack specificity^21^. As liver diseases are chronic, evolutive pathologies, patients would benefit from portable monitoring devices that enable rapid and simple assessment of liver function with high sensitivity and specificity.

An alternative biomarker of liver dysfunction is the presence of bile salts in serum. Bile salts are key components of bile which are critical for digestion in which they help absorption of fat^22^. Interestingly, serum bile salts have emerged as a general biomarker of liver disease, and are the gold-standard diagnostic method for pregnancy cholangitis^24,25^. In addition, several studies have pointed to bile salts as a general biomarker of interest for early cirrhosis, hepatitis, drug-induced liver injury. In addition, bile salts provide a highly specific and dynamic assessment of liver function. For example, serum bile salts are highly correlated with the obstruction state of the bile duct and rapidly decrease when biliary stenting is performed^26^. Furthermore, specific bile salt profiles may be associated with particular liver diseases^27–29^. Bile salts thus represent an ideal and specific biomarker for diagnosis and dynamic monitoring of liver disease. Yet, as for other biomarkers, current detection methods for bile salts based on enzymatic assays^30^ are only performed in a centralized fashion and cannot discriminate between different bile salts.

To address these challenges, we took advantage of the natural capacity of enteropathogenic bacteria to detect bile salts upon arrival into the gut to activate their virulence pathways^31,32^. We rewired sensing modules for bile salt detection into our modular bacterial receptor platform termed EMeRALD (**E**ngineered **M**odularized **R**eceptors **A**ctivated via **L**igand-induced **D**imerization) operating in *Escherichia coli*^*33*^. As the synthetic receptor operates in a surrogate, non-pathogenic host, we performed directed evolution of the sensing module and improved its limit-of-detection (LOD) and sensitivity. Finally, we optimized a colorimetric version of the system to operate in clinical samples. The resulting bactosensor can detect pathological levels of bile salts in serum from patients having undergone liver transplant and provides a signal detectable with the naked eye. The bile salt bactosensor provided here could support field monitoring of liver function for a wide range of conditions. Our work highlights the flexibility and modularity of the EMeRALD receptor platform for rapid characterization and engineering of novel sensing capabilities in whole-cell biosensors.

## RESULTS

### Engineering of synthetic bile-salt receptors in *E. coli* using the EMeRALD platform

Enteropathogenic bacteria such as *Vibrio cholerae* or *Vibrio parahaemolyticus* cause acute intestinal infections mediated by toxin secretion^34^. These pathogens use bile salts as an intestinal location signal to activate their virulence pathways. Bile salt sensing is mainly under the control of inner membrane sensor/cofactor couples, TcpP-TcpH for *V. cholerae*^*31*^ and VtrA-VtrC for *V. parahaemolyticus*^*35*^. As using pathogens as biosensors would involve significant host-specific regulation correction^35,36^ and biosafety containment issues, an alternative strategy is to rewire pathogen sensing modules of interest into a modular platform operating in a surrogate host (**Fig. 1a**).

**Figure 1.**
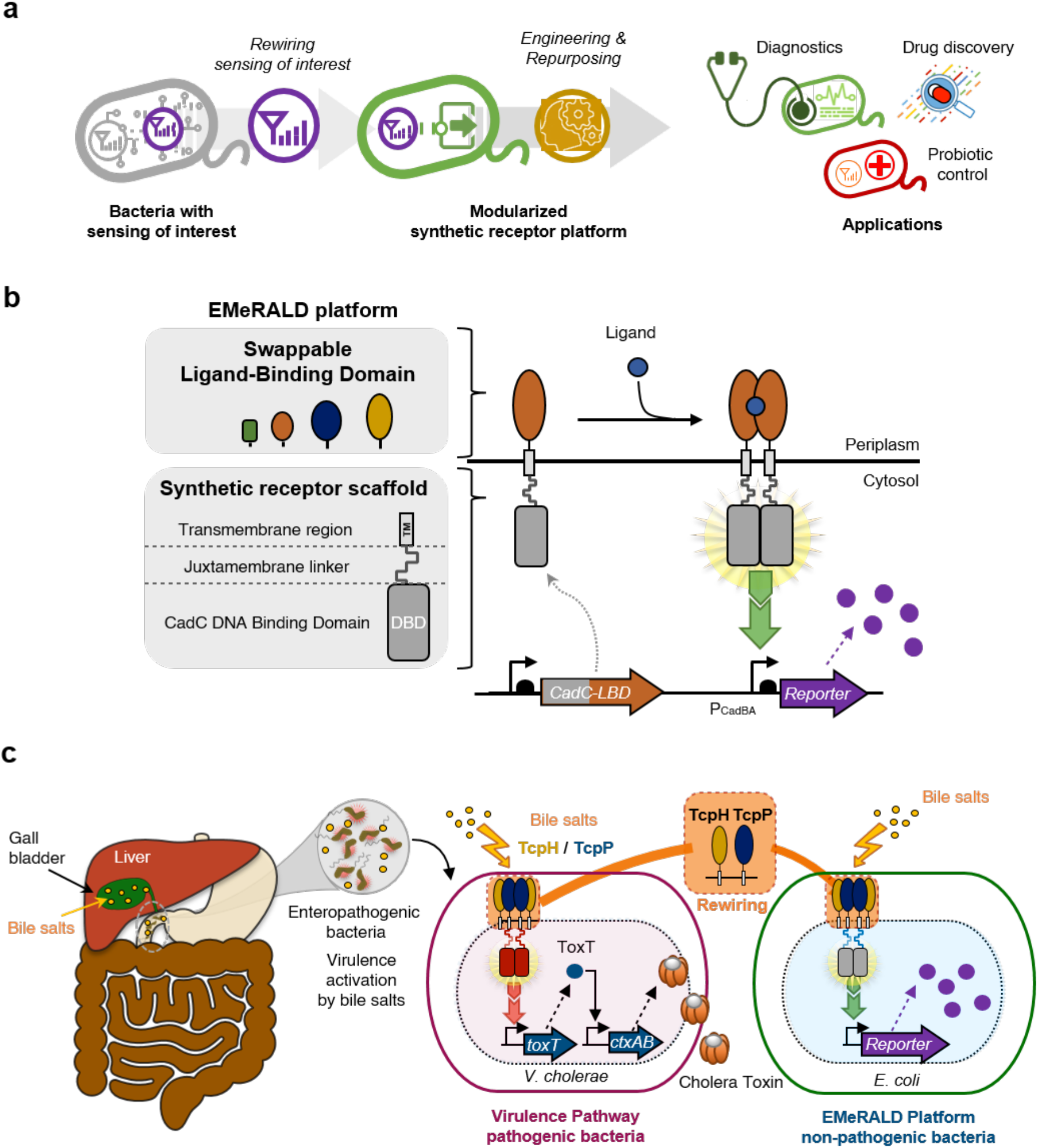
Design principles and architecture of EMeRALD-based bacterial sensors. **(a)** General strategy of rewiring bacterial sensing modules into surrogate hosts using a synthetic receptor platform. **(b)** Architecture and functional components of EMeRALD platform. The EMeRALD platform is composed of swappable Ligand-Binding Domains (LBD) that can be plugged into a synthetic receptor scaffold consisting of the DNA binding domain (DBD) of the CadC protein, a juxtamembrane (JM) linker, and a transmembrane region. The resulting synthetic transmembrane receptor is activated via ligand-induced dimerization and triggers reporter gene expression. **(c)** Rewiring bile salt sensing into *E. coli* using the EMeRALD platform. Enteropathogenic bacteria detect intestinal bile salts as host environmental cues for activating their virulence pathway. We plugged the *V. cholerae* bile salt receptor TcpP, and its cofactor protein TcpH, into the EMeRALD platform operating in the surrogate host *E. coli* to build a synthetic bile salt receptor controlling expression of a reporter gene.

The EMeRALD receptor platform, which we recently designed^33^ (**Fig. 1b**), is derived from membrane-bound one-component systems, which are bitopic proteins with a typical architecture of a cytoplasmic DNA Binding Domain (DBD), a juxtamembrane linker, a transmembrane region, and a periplasmic Ligand Binding Domain (LBD). Direct fusion between the LBD and the DBD provides a simple yet efficient solution to transduce incoming signals into a transcriptional output^37,38^. The EMeRALD platform operates in *E. coli* and uses the DBD from the CadC pH and lysine sensor which is inactive in its monomeric state^39^. Ligand-induced dimerization of the LBD triggers dimerisation of the cytoplasmic DBD and transcriptional activation^40^. This straightforward mechanism offers the potential to modularize receptor sensing and signaling by domain swapping. We previously built a synthetic receptor responding to caffeine by using a nanobody for this ligand as LBD^33^.

To engineer a chimeric bile-salt receptor in *E. coli*, we fused the *V. cholerae* TcpP bile-salt sensing module and its transmembrane region (TM) to the DNA binding domain of CadC (**Fig 1c**). As a reporter, we placed superfolder Green Fluorescent Protein (sfGFP)^41^ under the control of the CadC target promoter, pCadBA. We also expressed the cofactor protein TcpH, previously described to protect TcpP from proteolysis by the *V. cholerae* RseP protease once dimerized in response to bile salts^42^. We first confirmed using inducible gene expression systems that co-expression of the TcpH cofactor was necessary for TcpP function, using the primary bile salt taurocholic acid (TCA) as a ligand (**Supplementary Fig. 1-3**). These data suggest that the RseP homolog present in *E. coli* (UniProt:P0AEH1) can degrade TcpP when not bound by TcpH. We also found that the relative expression level of CadC-TcpP and TcpH were critical parameters affecting system performance (**Supplementary Fig. 3**).

We then placed both proteins under the control of constitutive promoters^43^. We used the strong constitutive promoter P5 for TcpH and three different constitutive promoters of increasing strengths, P9, P10, and P14, for CadC-TcpP (**Fig. 2a**) and tested their response to TCA^42^ (**Fig. 2b, Supplementary Fig. 4-5**). We found that the P9-CadC-TcpP variant had the lowest LOD, highest dynamic range, and highest signal strength. These data confirm that the stoichiometry between CadC-TcpP and TcpH is a key parameter influencing receptor performance.

**Figure 2.**
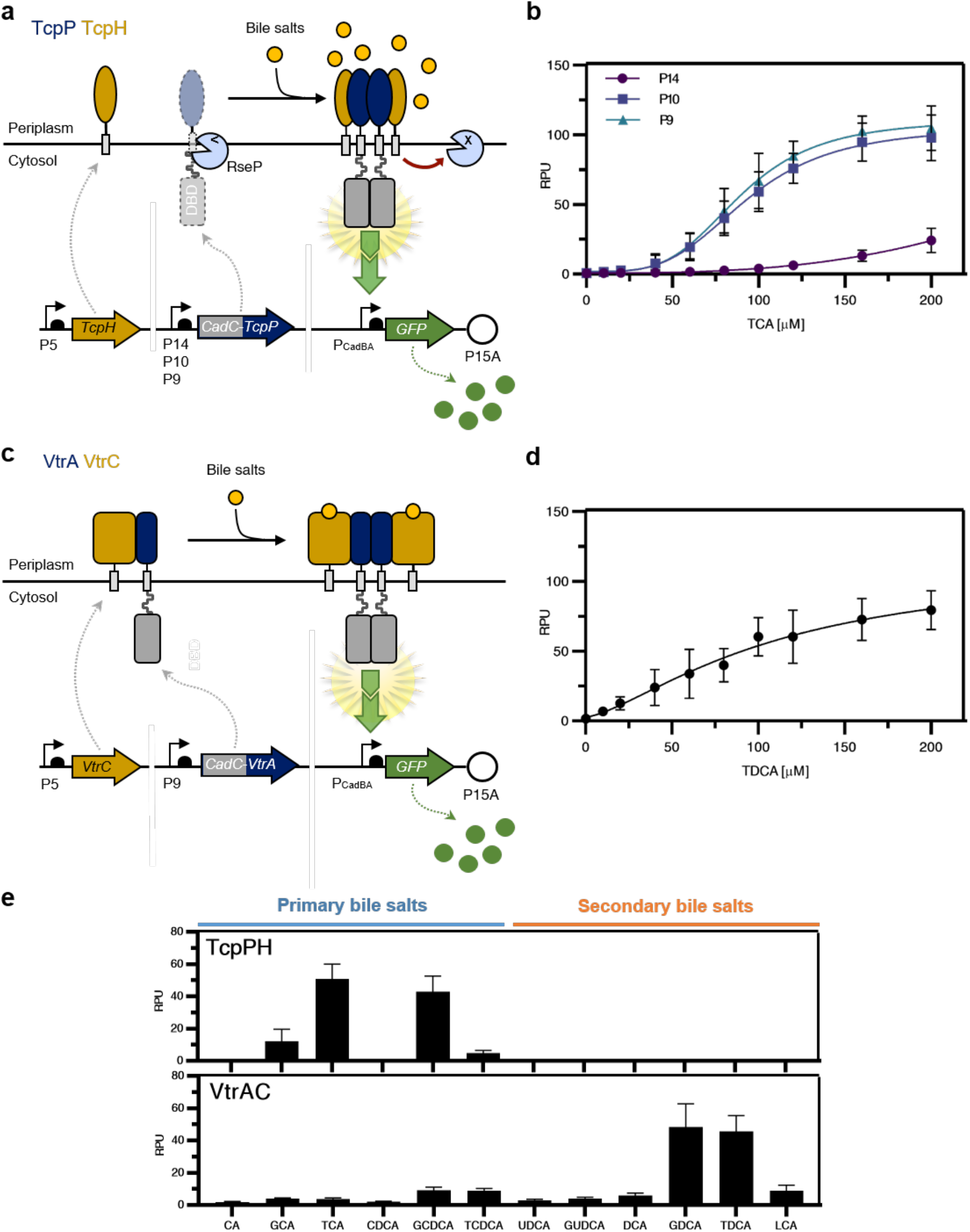
Design, implementation and characterization of synthetic bile salt sensors. **(a)** Overview of the TcpPH-EMeRALD system. The CadC DNA binding-domain (DBD) is fused to the transmembrane and periplasmic domains of TcpP. Three constitutive promoters (P14, P10, and P9) were tested to tune the transcription level of CadC-TcpP. Transcription of the TcpH cofactor is under the control of the constitutive promoter P5. In the absence of bile salts, CadC-TcpP is probably degraded by an endogenous *E. coli* homolog of the *V. cholerae* protease RseP. In the presence of bile salts, CadC-TcpP dimerizes and forms a stable complex with TcpH that protects it from proteolysis. The CadC-TcpP dimer then activates downstream expression of the GFP reporter. **(b)** Transfer function of TcpPH-EMeRALD receptors controlled by different promoters in response to increasing concentrations of the bile salt taurocholic acid (TCA). **(c)** Overview of the VtrAC-EMeRALD system. The CadC DBD is fused to the transmembrane and periplasmic domains of VtrA. CadC-VtrA and VtrC are under the control of the P9 and P5 promoters, respectively. Bile salts binding to VtrA/VtrC heterodimeric complexes promote oligomerization of CadC-VtrA and activate downstream expression of the GFP reporter. **(d)** Transfer function of VtrAC-EMeRALD receptor to increasing concentrations of the bile salt taurodeoxycholic acid (TDCA). **(e)** Bile salt specificity profiles for TcpPH-EMeRALD and VtrAC-EMeRALD systems. The full names and molecular structure of the different bile salts are listed in Supplementary Fig.1. All experiments are the mean of three experiments performed in triplicate on three different days. Error bars: +/- SD. RPU: reference promoter units.

We then assessed the versatility of the EMeRALD platform by connecting the VtrA/VtrC sensor system from *V. parahaemolyticus*^*32*^ (**Fig. 2c**). We built a dual-expression system consisting of P9-CadC-VtrA and P5-VtrC, and tested its response to its canonical ligand taurodeoxycholic acid (TDCA). We found that the VtrA/VtrC EMeRALD system was functional with a slightly higher LOD, similar dynamic range and signal strength than the TcpP/TcpH EMeRALD system (**Fig. 2d, Supplementary Fig. 6**). These results highlight the modularity and scalability of the EMeRALD platform, which supports the connection of different sensing modules to the receptor scaffold.

### Synthetic bile salt receptors exhibit different specificity profiles

We then assessed the specificity profile of the synthetic bile salt receptors. Bile salts are classified in two categories: primary bile salts (including taurocholic acid) are produced by the liver while secondary bile salts arise from modification of primary bile salts by gut microbiome metabolism. Primary bile salts are upregulated in serum and urine of patients with liver disease^27–29^. Previously identified virulence activating factors for *V. cholerae* include taurocholic acid, glycocholate, and cholic acid^42^. We measured the response of the bactosensor to a panel of twelve different bile salts, including both primary and secondary types (**Fig. 2e, Supplementary Fig. 7-8**). Interestingly, the CadC-TcpP system was highly specific for primary conjugated bile salts (especially TCA and GCDCA), while not responding to secondary bile salts. On the other hand, the CadC-VtrA system has a larger spectrum of bile salt specificity, mainly responding to secondary conjugated bile salts GDCA and TDCA. Due to the link between primary bile salts and liver diseases, we selected the TcpP/TcpH system to develop a bile salt bactosensor for medical diagnosis.

### Directed evolution of TcpP sensing module for improved LOD and higher sensitivity

Sensor sensitivity and LOD are key parameters for biosensors applications. We aimed to identify key residues determining the sensitivity of the TcpP sensing module, and targeted those to improve synthetic receptor sensitivity and LOD. To do so, we coupled comprehensive mutagenesis with functional screening and Next-Generation Sequencing (NGS), an approach which supports the identification of functional variants together with the sequence determinants within local structural motifs^44–46^. This strategy has also been used to engineer orthogonal two-component systems^47^.

Transition from intramolecular to intermolecular disulfide bonds between TcpP monomers is a key determinant of TcpP response to bile salts and is mediated by two cysteine residues, Cys207 and Cys218^42^. By performing multiple sequence alignments of different TcpP bacterial homologs (**Supplementary Fig. 9**), we found a significant conservation of the amino-acids flanked by these two cysteines (**Fig. 3a**). Secondary structure prediction and *ab initio* 3D prediction using the Rosetta modelling suite (**Fig. 3b and Supplementary Fig. 10**) suggested that each cysteine was located in rigid beta-sheets separated by a flexible loop region between Asn211 and Gln214. This loop propensity to form a turn would allow the two beta sheets and the cysteines to come in close proximity and form an intramolecular disulfide bond. We hypothesized that the flexibility of the turn region between Cys207 and Cys218 was a key parameter controlling the transition rate between the two states, and that altering its amino-acid composition could change the system’s sensitivity to bile salts.

**Figure 3.**
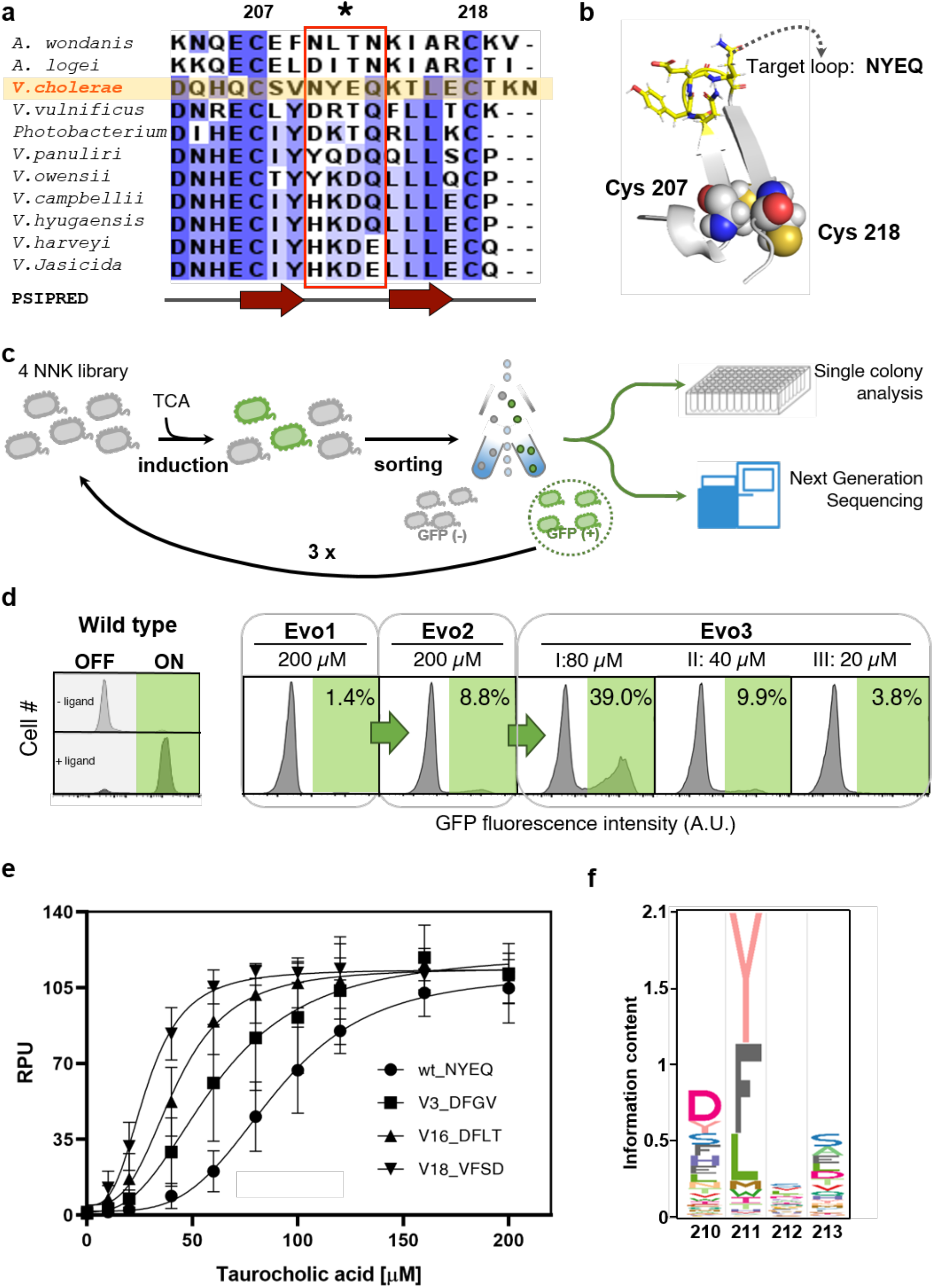
Directed evolution of TcpP for improved LOD and sensitivity. **(a)** Multiple sequence alignment of the C-terminal periplasmic domains from TcpP homologous proteins. Colors indicate sequence identities. Secondary structures prediction by PSIPRED is shown below. The red dashed box indicates the region of interest for mutational scanning. **(b)** *Ab initio* modeling of the C-terminal region of the TcpP sensing module. The two cysteine residues are shown as spheres. The amino acid residues NYEQ in the loop region of interest are labeled as sticks. **(c)** Schematic diagram of the screening procedure to obtain functional TcpP loop variants. The cell library was submitted to several rounds of sorting based on fluorescent signal output produced in response to bile salts. Individual clones were then isolated and characterized while the sequence of the mutated loop for the whole pool of isolated variants was analyzed by NGS. **(d)** Screening conditions and results of TcpP directed evolution. The gate used for cell sorting is colored in green. The x-axis indicates the fluorescence intensity in arbitrary units (A.U.). The gate for fluorescence-activated cell sorting is determined using the fluorescence distribution of TcpPH-EMeRALD sensor with or without the 200 μM TCA (left panel). **(e)** Transfer function of TcpP functional variants with improved sensitivity in response to increasing concentrations of TCA. Amino acid sequences of the loop regions are indicated. **(f)** Sequence logos of the flexible loop region from selected TcpP functional variants derived from NGS data. For (e) experiments are the mean of three experiments performed in triplicate on three different days. Error bars: +/- SD.

We thus built a comprehensive mutational library (NNK x 4, theoretical library complexity ≌ 1.05 × 10^6^ variants, see methods for details) targeting the NYEK residues inside the turn, and cloned it into the plasmid constitutively expressing CadC-TcpP and producing GFP in response to bile salts (**Fig. 3c**). The resulting library was induced with TCA, and GFP-positive variants were isolated by fluorescence-activated cell sorting (FACS). We performed three rounds of enrichment (200*μ*M of TCA as ligand in 1st and 2nd rounds of selection, and 20 µM for the 3rd round) and observed an increasing fraction of the cell population responding to different ligand concentrations (20 to 80 µM) (**Fig. 3d**). We collected, cultured, and sequenced single variants and tested their response to TCA (**Fig. 3e**). We found that comprehensive mutagenesis of residues Asn210 to Gln213 could alter the limit of detection, the sensitivity, and the fold activation of our biosensor. The 3.3-fold difference in LOD between V18 and V22 (EC_50_ from 28.3 to 92.5 µM, **Table 1**) indicated the broad range of sensitivity engineering obtained by mutating the loop region of the TcpP sensing module. Further kinetic analysis revealed that the sequence variation of this loop region changes reaction speed and system interaction of bile salts with the synthetic receptor (**Supplementary Fig. 11**).

**Table 1.** Functional analysis of selected TcpP functional variants.

| # Variant | Sequence | EC50 (μM) | Response<br>EC50 (RPU) | Max fold<br>change |
| --- | --- | --- | --- | --- |
| V22 | YYVL | 92.531 | 56.19 | 69.06 |
| TcpPwt | NYEQ | 89.405 | 56.5 | 164.74 |
| V7 | WYVH | 86.029 | 53.37 | 51.31 |
| V78 | FYES | 84.668 | 59.62 | 69.06 |
| V11 | YYIV | 82.167 | 53.83 | 31.76 |
| V14 | TFLA | 81.636 | 62.97 | 222.67 |
| V3 | DFGV | 61.48 | 61.18 | 142.21 |
| V19 | FFKA | 59.186 | 61.29 | 127.75 |
| V16 | DFLT | 42.058 | 59.34 | 75.73 |
| V18 | VFSD | 28.344 | 58.99 | 84.92 |

To better understand the sequence features influencing the response of TcpP to bile salts, we sequenced the whole pool of enriched variants by next-generation sequencing (NGS). Surprisingly, the sequence features of functional variants were different from those expected from natural TcpP homologs (**Fig. 3f and Supplementary Fig. 12**). First, and in contrast with *wt* TcpP homologs, we observed a strong depletion of long-chain, negatively charged amino-acids (Asp and Glu) along with long-chain polar amino-acids (Asn and Gln) at position 211. Lysine at position 211 also appeared to be depleted in functional variants (despite being commonly found at this position in other TcpP homologous proteins). Second, amino acids with bulky aromatic side chain such as Phe and Tyr were highly conserved in selected functional variants, but not among TcpP homologs (with the notable exception of Tyr211 in *V. cholerae* TcpP), strongly indicating the important role of aromatic residues at position 211 in the function of *V. cholerae* TcpP. We chose the best engineered variant, termed TcpP18, for further development of a clinical bactosensor.

### Development of a colorimetric version of the bactosensor

Colorimetric assay provides a simple and intuitive method for simple and direct estimation of test results by the naked eye. In addition, colorimetric assays support straightforward development of quantitative assays using smartphone-based platforms for POC or home-based diagnosis^48^. We used TcpP18 coupled with the reporter beta-galactosidase LacZ (termed TcpP18-LacZ) and its substrate chlorophenol red-β-D-galactopyranoside (CPRG) to provide a colorimetric output^49^ (**Fig. 4a**, see methods for details). Similarly to the biosensor equipped with a GFP output, the bile salt specificity profile of the TcpP18-LacZ system was slightly shifted from TCA to GCDCA (**Fig. 4b, Supplementary Fig. 13**). We thus evaluated the LOD and signal output threshold of TcpP18-LacZ in response to increasing concentrations of GCDCA. We also explored the influence of varying cell density and incubation time (**Fig. 4c** and **Supplementary Fig. 14**). Increasing cell density or incubation time both improved the dynamic and operating ranges of TcpP18-LacZ; however, background signal also increased. After optimization, the Tcp18-LacZ was able to detect clinically relevant bile salt concentrations, such as those corresponding to early cirrhosis (~ 5.2 μM GCDCA)^50^ or nonalcoholic fatty liver diseases with hepatocellular carcinoma (~ 56 μM GCDCA)^51^.

**Figure 4:**
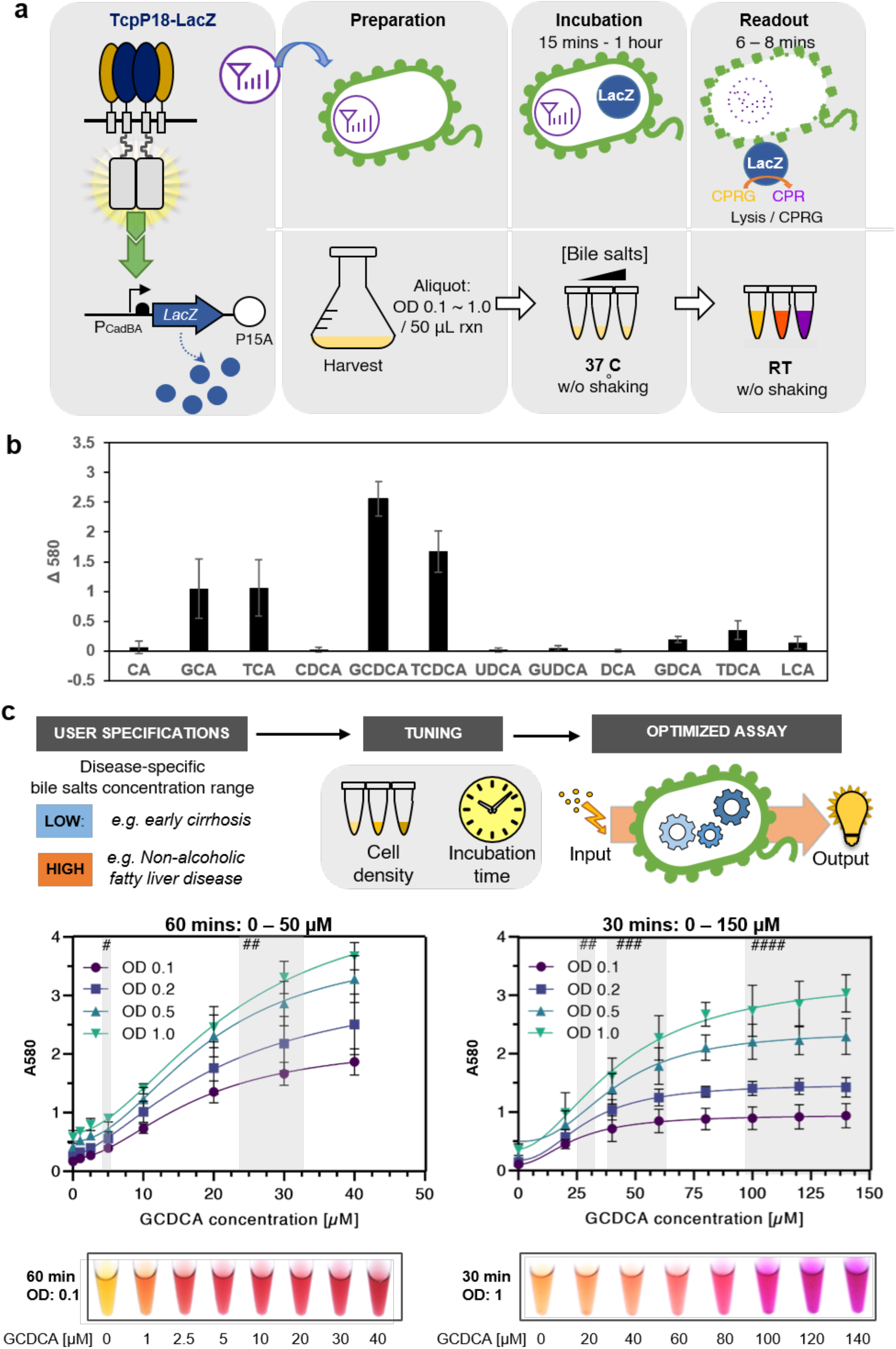
Colorimetric assay for bactosensor mediated bile salts detection. **(a)** Schematic diagram for the design and operation procedure of the TcpP18-LacZ system for bile salt detection. The TcpP18-LacZ uses the TcpP loop variant V18 as sensing module for lower LOD and higher sensitivity and LacZ as a colorimetric reporter, with CPRG as a substrate which is converted to chlorophenol red turning the reaction from yellow to purple. **(b)** Bile salt specificity profile of the TcpP variant V18 characterized using TcpP18-LacZ sensor. Response of TcpP18-LacZ was quantified as ΔA580 (the difference in absorbance at 580 nm (A580) with or without ligand bile salts). **(c)** Optimisation and quantification of TcpP18-LacZ response to the bile salt glycodeoxycholic acid (GCDCA). The LOD and response dynamic range of TcpP18-LacZ were fine-tuned by varying the cell concentration and incubation time. The grey bars with different (#) indicate serum GCDCA concentrations in patients serum with different stages of liver disease extracted from the litterature^26,50,51^ (#: early cirrhosis; ##: biliary obstruction; ###: non-alcoholic fatty liver disease (NASH) with hepatocellular carcinoma; ####: NASH with cirrhosis). All experiments are the mean of three experiments performed in triplicate on three different days. Error bars: +/- SD. See methods, main text and SI for details.

### Bactosensor-mediated detection of elevated bile salts levels in serum from patients with liver transplant

We then prototyped our bactosensor for the detection of bile salts in clinical conditions. To do so, we tested the sensor on samples from patients having undergone liver transplantation. After liver transplant, the main complications are bile ducts stenosis and acute cellular rejection. In order to detect these complications at an early stage, liver tests are performed regularly. Serum bile salt concentration has been shown to be a good indicator for the assessment of liver dysfunction after liver transplantation^52^. A field-deployable method for bile salt assessment would greatly improve the monitoring of these patients, ultimately allowing fine grained monitoring performed at-home by the patients themselves.

We tested our bactosensor in clinical 21 serum samples from liver transplantation patients (**Fig. 5a and Supplementary Fig. 15**). The patients were followed at the Montpellier hospital after their liver transplant, most of them having been performed in the last 2 years **(see Supplementary Information)**. These patients had received a liver transplant for end-stage liver disease as a result of alcoholic related liver disease or non-alcoholic fatty liver disease, chronic cholangitis or liver cancer. A complete hepatic check-up was performed, and serum bile salts were also measured (**Supplementary Tables 3-5** for clinical data). We found that patients who had a high potential of acute cellular rejection (ACR) after liver transplantation (serum bile acid > 37 μM)^53^ had significant and visible colorimetric signal changes (**Supplementary Fig. 16**) in bactosensor assays. Three patients in particular raised our attention: patients #5, 10 and 13. These three patients had elevated serum bile salts concentration. Two of them (5 and 10) presented abnormalities in their hepatic enzymatic values (ASAT, ALAT, GGT, Pal, and bilirubin). For these patients, the bile salt bactosensor produced the strongest colorimetric change easily detectable with the naked eye (**Fig. 5b**). We further compared the results of the bactosensor with a commercial bile salt enzymatic assay (**Fig. 5c**). The high correlation between enzymatic serum bile salt assay and our bactosensor measurements (r = 0.908, p < 0.0001) indicates the reliability of our assay. In addition, we compared the results from the bactosensor with other serum liver biomarkers (**Supplementary Fig. 17**), and found a high correlation to a common liver biomarker, total bilirubin. These results indicate that our bactosensor is able to provide a simple, reliable, and cost-effective method for monitoring patient condition after liver transplantation.

**Figure 5:**
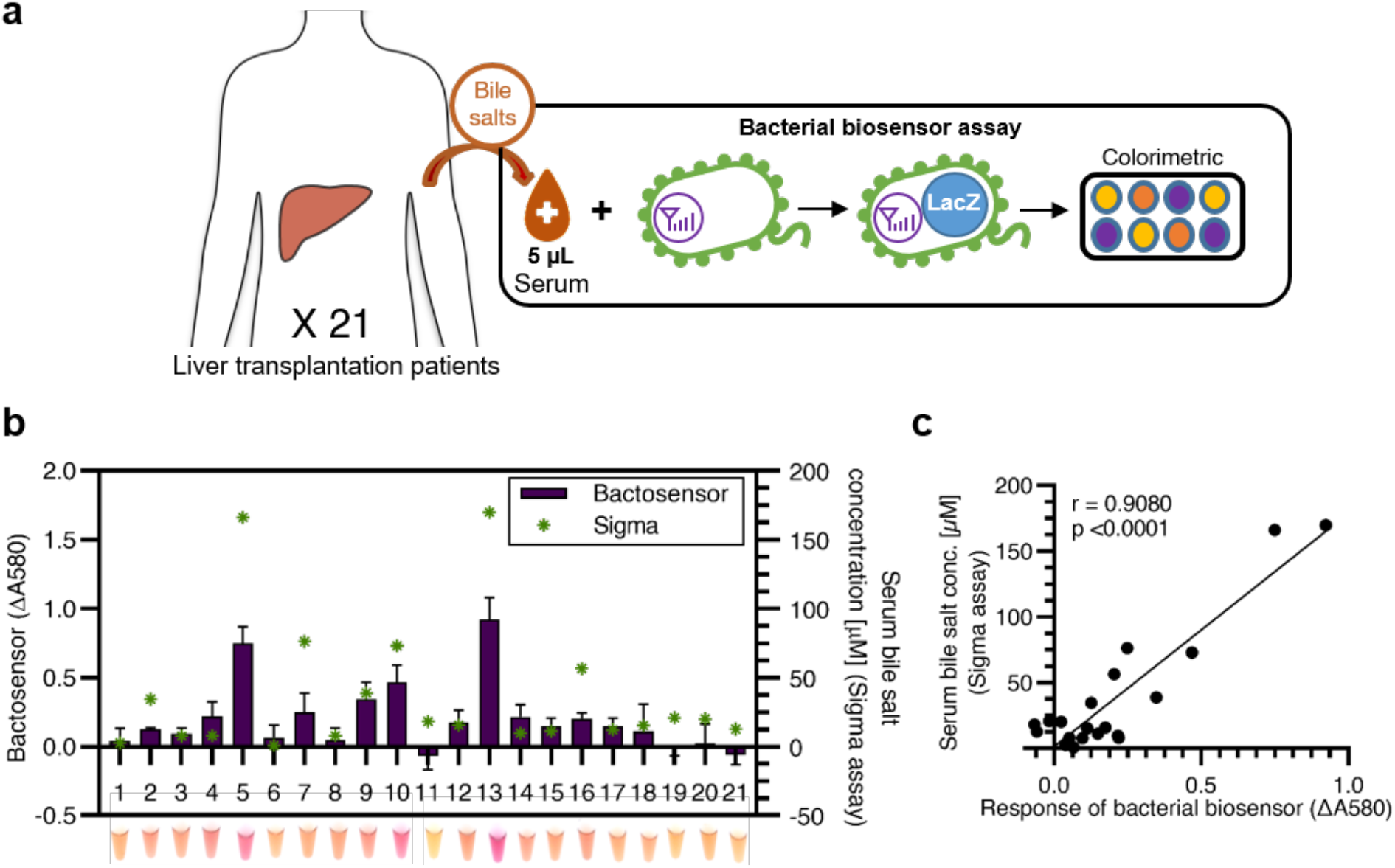
Bactosensor-based pathological bile salt detection in clinical samples. **(a)** Serum samples from 21 liver transplantation patients being followed at a Montpellier hospital were analyzed using a bactosensor equipped with the TcpP18-LacZ sensor as described in **Figure 4a**, with a 10-fold dilution and 2 hour incubation time (**Supplementary Fig. 14**). **(b)** Comparing the analysis result of 21 clinical serum samples between TcpP18-EMeRALD-LacZ and a bile salt assay kit. The response of TcpP18-LacZ is shown in black bars, left axis. The serum total bile salt concentration measured by a bile salt enzymatic assay kit is labeled in green asterisks, right axis. **(c)** Correlation between serum bile salt detection results from TcpP18-LacZ sensor and bile salt concentration measured by enzymatic assay. All experiments are the mean of three experiments performed in triplicate on three different days. Error bars: +/- SD. See methods, main text and SI for details.

## DISCUSSION

Microbes detect and process myriad environmental signals, providing a vast sensing repertoire for engineering biosensors usable for several applications. However, the intricacy of signaling networks and our limited understanding of their biochemical properties restrain their direct use. Here we presented a general strategy to rewire sensing modules of interest into a well-characterized synthetic receptor platform, EMeRALD, enabling their fine tuning and repurposing for novel biomedical applications. In the future, these receptors could be coupled with genetic circuits performing multiplexing logic, memory and signal amplification to engineer even more sophisticated whole-cell biosensors^54,55^.

We were able to connect two different bile salt sensing modules having different specificity profiles, the TcpP/H and VtrA/C systems, which respond mostly to primary and secondary bile salts, respectively. Importantly, our capacity to engineer synthetic bile salt signaling using only these protein domains demonstrates that these modules are the only essential components required for bile salt sensing in their natural host. By performing directed evolution of TcpP, we improved the sensitivity and decreased the LOD of the sensor. In addition, we discovered previously unknown amino-acid sequence features influencing TcpP function and potentially relevant for *V. cholerae* virulence. For instance, we found a stringent functional requirement for the presence of Tyr211 residue in the C-terminal loop of *V. cholerae* TcpP. The requirement for amino acids with a nonpolar aromatic side chain at Tyr211 indicates rigorous steric interactions located in the C-terminal loop region of *V. cholerae* TcpP. The exact role of Tyr211 is still unclear, but this residue may affect TcpP-TcpP dimer formation or TcpP-TcpH interaction. While the unique presence of Tyr at the 211 position in *V. cholerae* was visible on multiple sequence alignments, the link between this residue and TcpP function could not be inferred by this approach.

This discovery exemplifies how our synthetic receptor platform offers a powerful strategy to interrogate the sequence-function relationship of bacterial sensing modules in a massively-parallel fashion. Such studies in the natural pathogenic host would have been tedious because of complex pathway regulations and safety issues would have limited the final bactosensors to a few expert groups. In contrast, our platform provides a straightforward and scalable strategy to study pathogen signaling in surrogate hosts, usable by the larger scientific community. The EmeRALD system, coupled with the deep-mutational scanning, methods to navigate protein sequence space^56–59^, designer libraries, and Flow-Seq analysis^46^ can serve as a general platform for studying bacterial sensing modules, engineering their ligand specificity and their response properties. In particular, our system is ideal for understanding transmembrane one-component signaling, which mechanism and regulation are currently underexplored^38^. On the same line, EMeRALD receptors could be applied to the discovery of inhibitors of virulence pathways through screening of chemical or natural substances libraries.

Here we show the application of EMeRALD to the field of medical diagnostics by detecting abnormal bile salt levels in patient samples. As bile salts dysregulation and gut dysbiosis are critical in the pathogenesis of liver diseases or gastrointestinal cancer^60–65^, there is an urgent need for POC assays for bile salt monitoring^66^. Our bactosensor results correlate well with hospital tests and produce a very strong signal detectable with the naked eye for the three most critical patients, demonstrating the potential of our technology for rapid POC estimation of liver dysfunction. In addition, our system uses very small sample volumes (5 μL) providing an efficient method for POC diagnostics. Yet, given the size of our patient cohort and its diversity, further clinical studies are needed to move our method towards the clinic. Interestingly, the bile salt sensing system could also be applied to control the activity of engineered probiotics upon arrival in the gut, as recently proposed in *Bacteroides thetaiotomicron*^*67*^.

The EMeRALD platform can accommodate natural or synthetic^33^ sensing modules, providing a versatile and scalable solution to develop new sensing modalities in bacteria. We anticipate that this platform will be repurposed for other applications in medical diagnostics, bacterial therapeutics, and environmental monitoring, supporting the transition of bacterial biosensors towards real-world applications.

## METHODS

### Plasmids and strains

Genetic parts of constructs used in this study are provided in Supporting Information. All constructs were cloned into plasmid J64100_p15A with p15a origin of replication and chloramphenicol resistance gene by isothermal Gibson assembly. All experiments were performed using *E. coli* strain NEB10β (New England Biolabs). Plasmids and materials will be made available through Addgene. Sequences will be deposited into GENBANK.

### Functional characterization of synthetic bile salt receptors

Plasmids encoding different constructs were transformed into chemically competent *E. coli* NEB10β (New England Biolabs), plated on LB agar plates supplemented with 25 μg/mL chloramphenicol and incubated at 37°C overnight. For each measurement, three fresh colonies were picked and inoculated into 5 mL of LB/chloramphenicol and grown at 37°C with vigorous shaking for 16-18 hours. In the next day, the cultures were diluted 1:100 into 1 mL of LB/chloramphenicol medium with different concentration of bile salts in 96 deep-well plates (Greiner bio-one), incubated at 37°C with vigorous shaking for a further 4 hours and analyzed by flow cytometry. All experiments were performed at least 3 times in triplicate on three different days. For bile salt specificity profiles for TcpPH-EMeRALD and VtrAC-EMeRALD systems, experiments were performed using each bile salt at a 80µM concentration. All chemicals used in this research were purchased from Sigma-Aldrich.

### Calculation of Relative Promoter Units (RPUs)

Fluorescence intensity measurements among different experiments were converted into RPUs by normalizing them according to the fluorescence intensity of an *E. coli* strain NEB10β containing a reference construct and grown in parallel for each experiment. We used the constitutive promoter J23101 and RBS_B0032 as our in vivo reference standard and placed superfolder GFP as a reporter gene in plasmid pSB4K5. We quantified the geometric mean of fluorescence intensity (MFI) of the flow cytometry data and calculated RPUs according to the following equation:

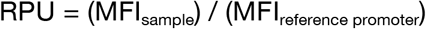

### Flow cytometry analysis

Flow cytometry was performed using an Attune NxT cytometer coupled with high-throughput autosampler (Thermo Fisher Scientific) and Attune NxT™ Version 2.7 Software and BD LSR Fortessa (Becton Dickinson), with FACSDiVa software. 30,000 cells were collected for each data point. Flow cytometry data were analyzed using FlowJo (Treestar Inc., Ashland, USA). All raw data values are in Supplementary Information. FCS files have been deposited in Flow Repository (https://flowrepository.org/). IDs are available in supplementary file 2.

### 4 × NNK library construction

Using P9-CadC-TcpP plasmid as template, the insert with 4 × NNK library was amplified by Phusion Flash High-Fidelity PCR Master Mix (Thermo Fisher Scientific) with the following primers: I5 - GGTCTCACAGCATCAATGTTCGGTGNNKNNK NNKNNKAAGACGTTGGAATGTACTAAGAACTAATAAAC, and I3 - GGTCTCAGGTCAATAATACCGAAACTATTTTATTGTTC; the vector was amplified with the following primers: V5:GGTCTCAGACCACTTCCGAGTAGAATCG, and V3: GGTCTCAGCTGGTCGATAGCAAAGGTCAAC. The purified fragments were first digested with BsaI-HFv2 (New England Biolabs), and then ligated by T4 ligase at 4°C overnight. 5 μg of ligation product was transformed into *E*.*coli* strain NEB10β with electroporation.

### Cell sorting

Cell sorting was performed using a S3 cell sorter (Bio-rad). 100,000 cells under different induction conditions (as shown in Figure 3) were gated and collected in SOC medium at each round of sorting. The sorted cells were further inoculated in 10 mL of LB/chloramphenicol medium at 37°C with vigorous shaking for a further 16-18 hours. The cell cultures positively selected cells were then applied to the next round of selection. For the first two rounds, the loop variant library was induced with 200 μM of TCA. In the third round of evolution, cells were further induced at different concentrations of TCA to isolate more sensitive variants. For each round of screening, cells from an overnight culture were diluted 1:100 in LB with or without TCA and grown for 16 hours at 37°C before being sorted.

### Rosetta modeling

Ab initio structural modeling. Structural models of the TcpP C-term segment (residues 182 to 211) were generated using the ROSETTA ab initio 3D prediction protocol ^68^ with 1D sequence and 2D predicted secondary structure as input data.

### Next Generation Sequencing

Cells sorted from the third round of evolution were further induced with 320 μM of TCA to collect the fully activated TcpP variants. Plasmids extracted from sorted cells were further amplified and added UMI barcodes with first round primers (see supplementary materials for sequence details). After PCR clean up, the first round PCR products were further amplified by 2nd round PCR primers with NGS index. The PCR products were further purified with AMPure XP paramagnetic beads to remove the contaminants. Samples were sequenced on Illumina PE250 platform at Novogen (Hong Kong, China) with paired-end reads of 250 bases.

### NGS data processing

The datasets generated and analyzed during the current study are available in the NCBI SRA repository (BioProject ID: PRJNA714981; https://www.ncbi.nlm.nih.gov/bioproject/714981). The python scripts for NGS data processing are available upon request. Sequence counts were converted into amino acid sequences and listed in supplementary file 1. Sequence logos were prepared by R package Logolas^69^.

### Colorimetric assay of bactosensors for bile salt detection

Overnight cultures of TcpP-LacZ were diluted in 1/250 fold into 100 mL of LB with 25 μg/mL chloramphenicol, incubated at 37 °C for 4 hours to reach an OD600 of around 0.4. The cells were put on ice for 30 mins and then spun down at 4°C and 2,200 g for 10 mins. The cell pellets were resuspended with LB medium (without antibiotics) at 1.1x of the final OD. 5 μL of different bile salts (prepared in LB medium) was added into 45 μL of resuspended cells in 96 well plates. The plate was incubated at 37°C for 10-60 mins without shaking. After incubation, 50 μL of B-PER solution (Thermo Fisher) with 800 μM CPRG was added into the cell culture directly. The mixture was further incubated for 6 mins at room temperature and then 50 μL of 1M sodium carbonate was added to stop the reaction. The absorbance at 580 nm was measured with a plate reader (Biotek; Cytation3). For the clinical sample assay, serum samples were heat-inactivated by incubation in a 56°C water bath for 30 mins. 5 μL of serum samples were added into 45 μL of resuspended cells in a 96 well plate and incubated at 37°C without shaking for 2 hours before lysis.

### Patient samples

Serum samples were consecutively collected from hospitalized or ambulatory liver transplant patients at the Hepatology and Liver Transplantation Unit, Hôpital Saint-Eloi in Montpellier (France) in the month of July 2020. All patients underwent liver transplantation between November 2006 and April 2020. Clinical and biological data were also recorded for every patient included in the study from their medical files. Abnormal liver enzymes were defined by : alanine transaminase (ALT) >41 U/L, aspartate transaminase (AST) >40 U/L, alkaline phosphatase (ALP) >130 U/L, gamma-glutamyl transferase (GGT) >60 U/l or total bilirubin (TBil) >21 µmol/L. The ethics committee of the University Hospital of Montpellier granted ethical approval (Number: 198711) and all patients signed an informed consent.

## Supporting information

Raw data values

NGS data

Flow repository IDs

## ACKNOWLEDGEMENTS

We thank members of the synthetic biology group and the CBS for fruitful discussions. We are grateful to the patients for participating in this study and providing their samples, and to the personnel of the Montpellier CHU hospital for collecting and preparing the samples. This work was supported by an ERC starting grant “COMPUCELL” to J.B. J.B. also acknowledges support from the INSERM Atip-Avenir program and the Bettencourt-Schueller Foundation. The CBS acknowledges support from the French Infrastructure for Integrated Structural Biology (FRISBI) ANR-10-INSB-05-01.

## AUTHOR CONTRIBUTIONS

H.J.C., A.Z., G.C. and J.B. designed experiments. H.J.C., A.Z. P.L.V., E.F.R. performed experiments. H.J.C., A.Z., P.L.V., E.F.R., G.C.,and J.B. analyzed experiments. I.C., L.M., M.M., and G.P.P. collected clinical samples and analyzed clinical data. J.G. performed bioinformatic analysis and structural modeling of TcpP. H.J.C., J.G., and M.C.G. analyzed the structural data. H.J.C. and J.B. wrote the paper. All authors participated in elaborating the final version of the manuscript and approved it.

