## Supplementary Materials for "A synthetic receptor platform enables rapid and portable monitoring of liver dysfunction via engineered bacteria"

### Rewiring bile salt sensing enables rapid and portable monitoring of liver dysfunction via engineered bacteria.R

**These supplementary materials contain:**

- Supplementary text.
- Supplementary figures S1 to S19.
- Supplementary tables 1 to 5.

**Supplementary text.**

**NGS primers used in this study**

**I. 1st round PCR for adaptor and UMI barcode:**

TcpP_Sensing_NGS_1st_Fw:
TCGTCGGCAGCGTCagatgtgtataagagacagNNNNNNNCCAGAAACTTAGCGAGCAGAAG

TcpP_Sensing_NGS_1st_Rv:
GTCTCGTGGGCTCGGagatgtgtataagagacagNNNNNNNgccacctgggatttccg

**II. 2nd round PCR for index insertion:**

| | |
|---|---|
| P5_bc1 | aatgatacggcgaccaccgagatctacacAACAACACtcgtcggcagcgtc |
| P5_bc2 | aatgatacggcgaccaccgagatctacacAACGGTGGtcgtcggcagcgtc |
| P5_bc3 | aatgatacggcgaccaccgagatctacacACCTTCTTtcgtcggcagcgtc |
| P5_bc4 | aatgatacggcgaccaccgagatctacacATTGAGCCtcgtcggcagcgtc |
| P5_bc5 | aatgatacggcgaccaccgagatctacacCCAGGAAGtcgtcggcagcgtc |
| P5_bc6 | aatgatacggcgaccaccgagatctacacCGTCCAATtcgtcggcagcgtc |

| | |
|---|---|
| P7_bc1 | caagcagaagacggcatacgagatCGTCCAATgtctcgtgggctcgg |
| P7_bc2 | caagcagaagacggcatacgagatGTGTCGTTgtctcgtgggctcgg |
| P7_bc3 | caagcagaagacggcatacgagatAACAACACgtctcgtgggctcgg |
| P7_bc4 | caagcagaagacggcatacgagatACCTTCTTgtctcgtgggctcgg |
| P7_bc5 | caagcagaagacggcatacgagatCCAGGAAGgtctcgtgggctcgg |
| P7_bc6 | caagcagaagacggcatacgagatAACGGTGGgtctcgtgggctcgg |

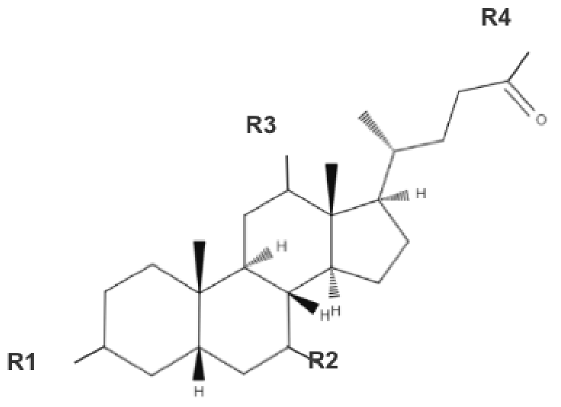

| Name | Abbreviation | R1 | R2 | R3 | R4 |
|---|---|---|---|---|---|
| Cholic acid | CA | OH | OH | OH | OH |
| Glycocholic acid | GCA | OH | OH | OH | $NHCH_2COO^-$ |
| Taurocholic acid | TCA | OH | OH | OH | $NHCH_2CH_2SO_3^-$ |
| Chenodeoxycholic acid | CDCA | OH(α) | OH(α) | H | OH |
| Glycochenodeoxycholic acid | GCDCA | OH(α) | OH(α) | H | $NHCH_2COO^-$ |
| Taurochenodeoxycholic acid | TCDCA | OH(α) | OH(α) | H | $NHCH_2CH_2SO_3^-$ |
| Ursodeoxycholic acid | UDCA | OH(α) | OH(β) | H | OH |
| Glycoursodeoxycholic acid | GUDCA | OH(α) | OH(β) | H | $NHCH_2COO^-$ |
| Deoxycholic acid | DCA | OH | H | OH | OH |
| Glycodeoxycholic acid | GDCA | OH | H | OH | $NHCH_2COO^-$ |
| Taurodeoxycholic acid | TDCA | OH | H | OH | $NHCH_2CH_2SO_3^-$ |
| Lithocholic acid | LCA | OH | H | H | OH |

**Supplementary Figure 1. Chemical structures of different bile salts and corresponding abbreviation used in this study.**

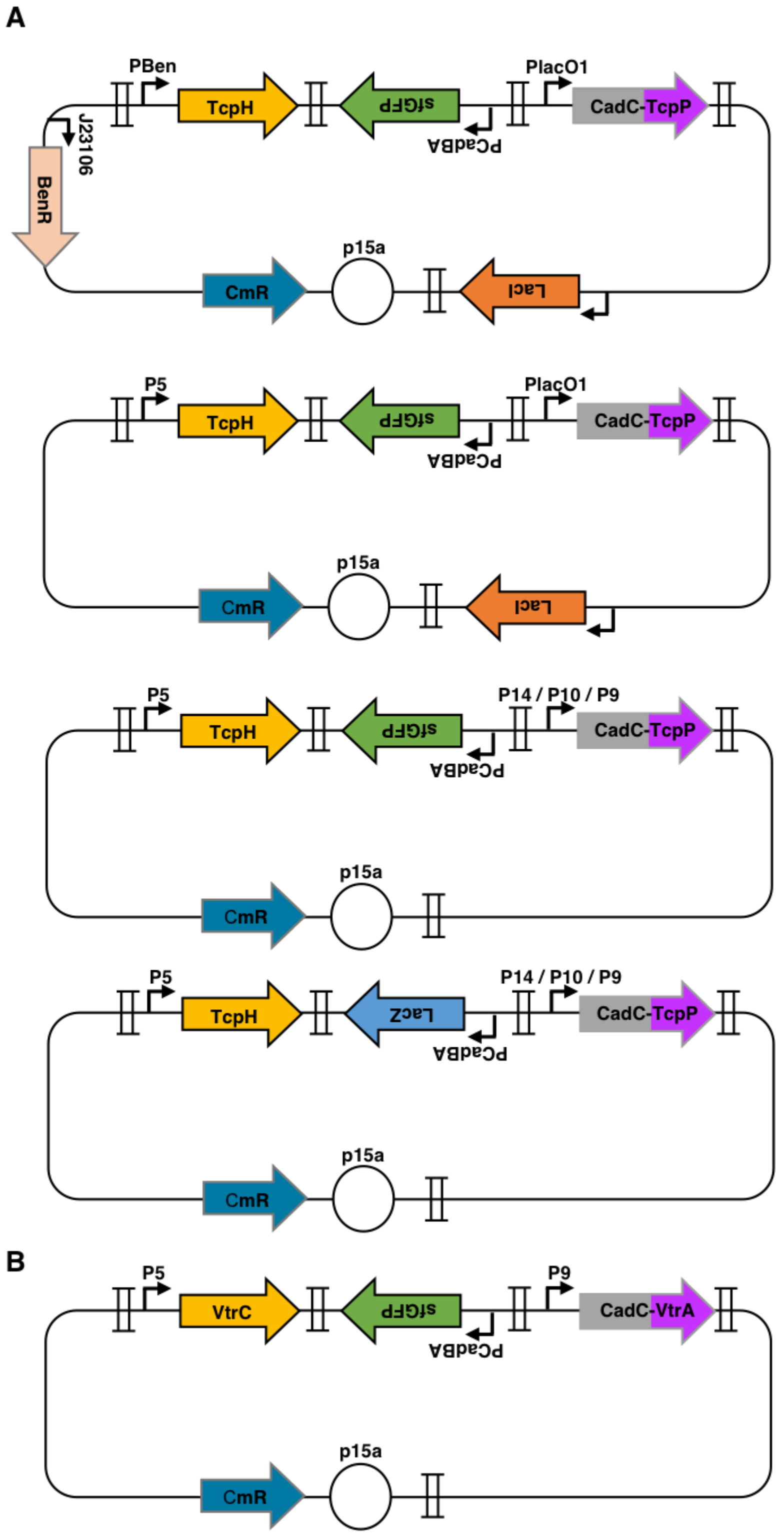

**Supplementary Figure 2: Plasmid maps of (A) inducible, mixed type and constitutive CadC-TcpP/TcpH system; and (B) constitutive CadC-VtrA/VtrC system.** *BenR*, the transcription factor BenR. *LacI*, the transcription factor LacI. CmR, the chloramphenicol resistance gene. Promoters are depicted by arrows; RBS are shown as solid half-circles; terminators are presented by II symbols.

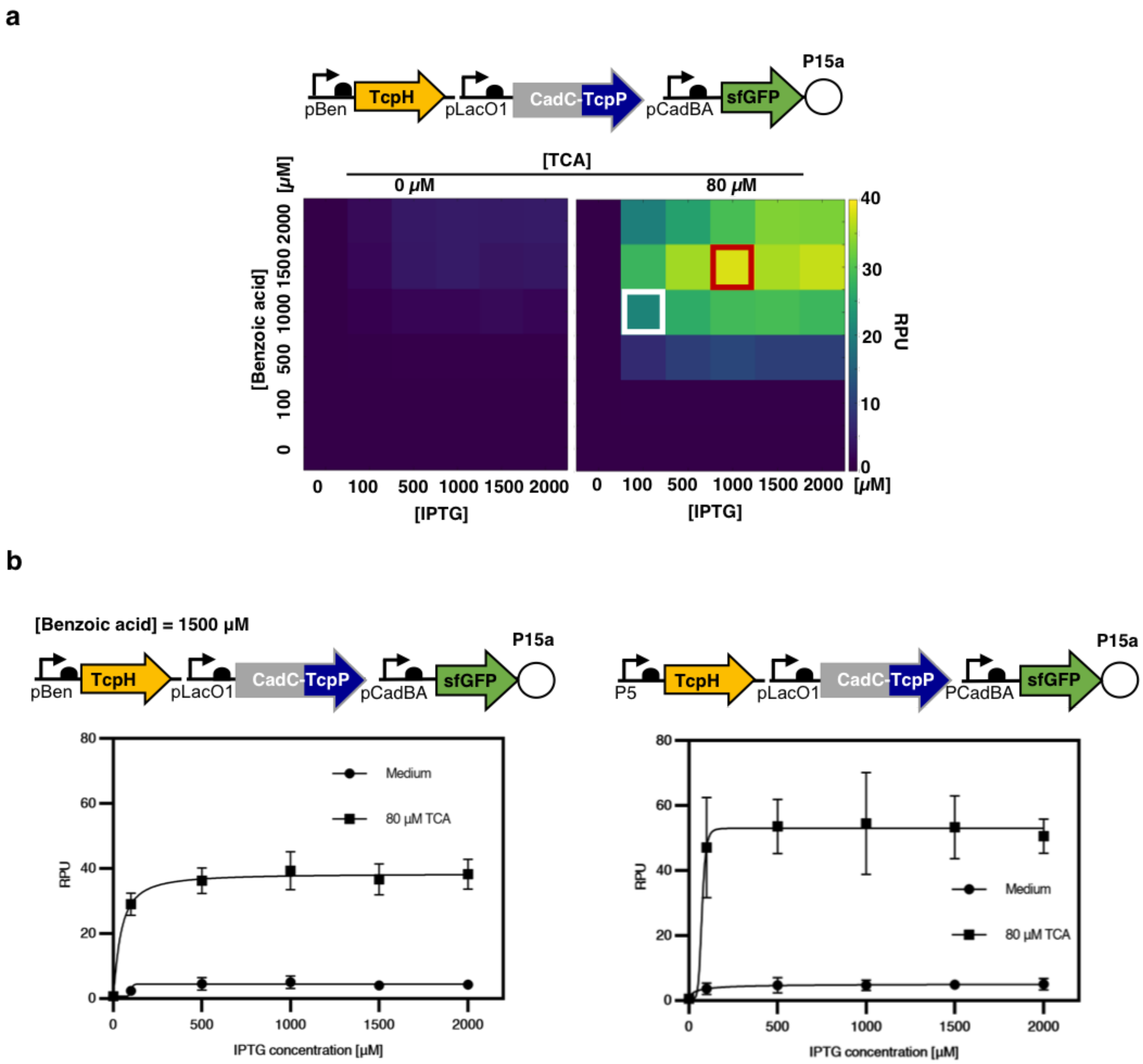

**Supplementary Figure 3: System performance of inducible CadC-TcpP/TcpH system.** (a) Response of inducible CadC-TcpP/TcpH system with or without the presence of ligand TCA at different expression levels induced by various concentrations of IPTG and benzoic acid. Schematic diagram of each genetic component and their corresponding control elements are listed above. RPU, reference promoter unit. The response with highest signal-to-noise ratio is highlighted with white square and the response with highest dynamic range is highlighted with red square. We found that there was an optimal expression ratio between the two proteins, i.e. maximizing protein expression did not lead to better output. (b) Comparing the different response between pLacO1-CadC-TcpP system with benzoic acid inducible (left panel) or constitutively expressed (right panel) TcpH, respectively. Inducible expression systems are useful to explore the parameter space of engineered systems. Yet the final bactosensor should operate using genes driven by optimal constitutive promoters. Constitutive expression reduces overall system complexity, DNA and metabolic footprint, and facilitates biosensor manipulation for real-world applications. Using constitutive promoters also helps avoiding toxicity or interference from the inducer molecule. Here we use the data from the inducible CadC-TcpP/TcpH system with highest dynamic range (induced by 1500 μM benzoic acid, Fig. S2a) to compare with the data from PlacO1-CadC-TcpP with constitutive expressed TcpH. We first placed the TcpH gene under the control of the strong constitutive promoter P5 [43], while keeping CadC-TcpP expression under IPTG control (right panel). The resulting combination showed a significant improvement in signal output level (swing) and signal-to-noise ratio compared to the system in which TcpH expression was under the control of the inducible pBEN promoter (left panel). These results suggested that the signal drop observed in Fig.S3a at high benzoic acid concentration might be due to a deleterious effect of the inducer benzoic acid on the cellular physiology.

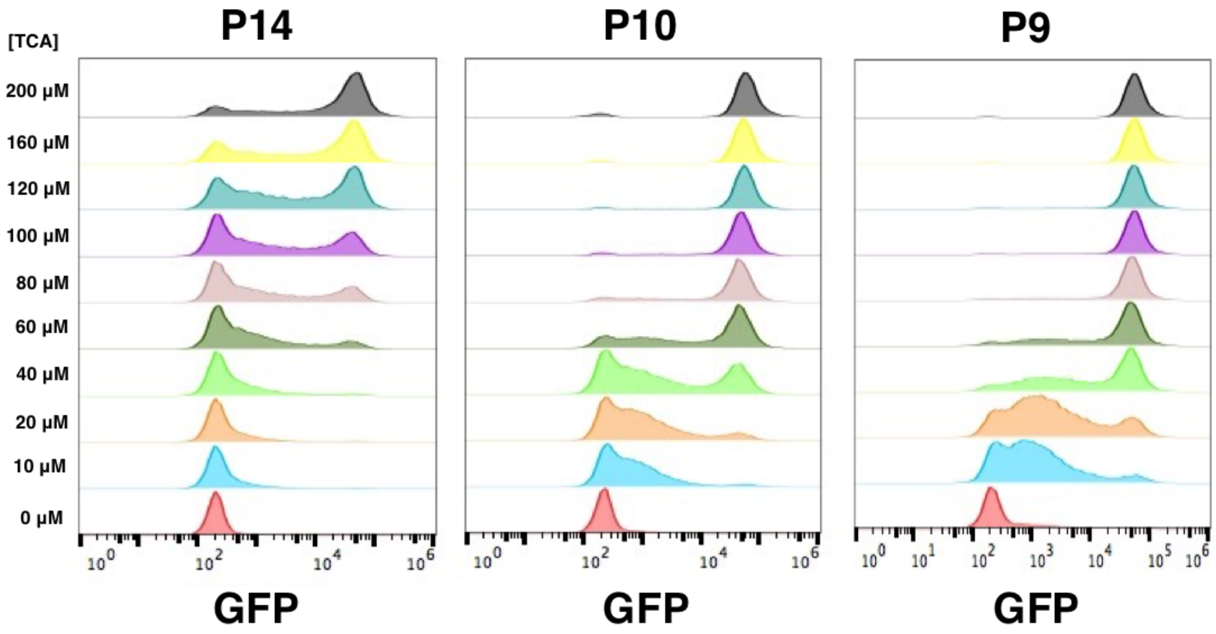

**Supplementary Figure 4: Response of CadC-TcpP promoter variants (with P5_TcpH) to increasing concentration of ligand TCA.** The TCA concentration is labeled at the left-y-axis.

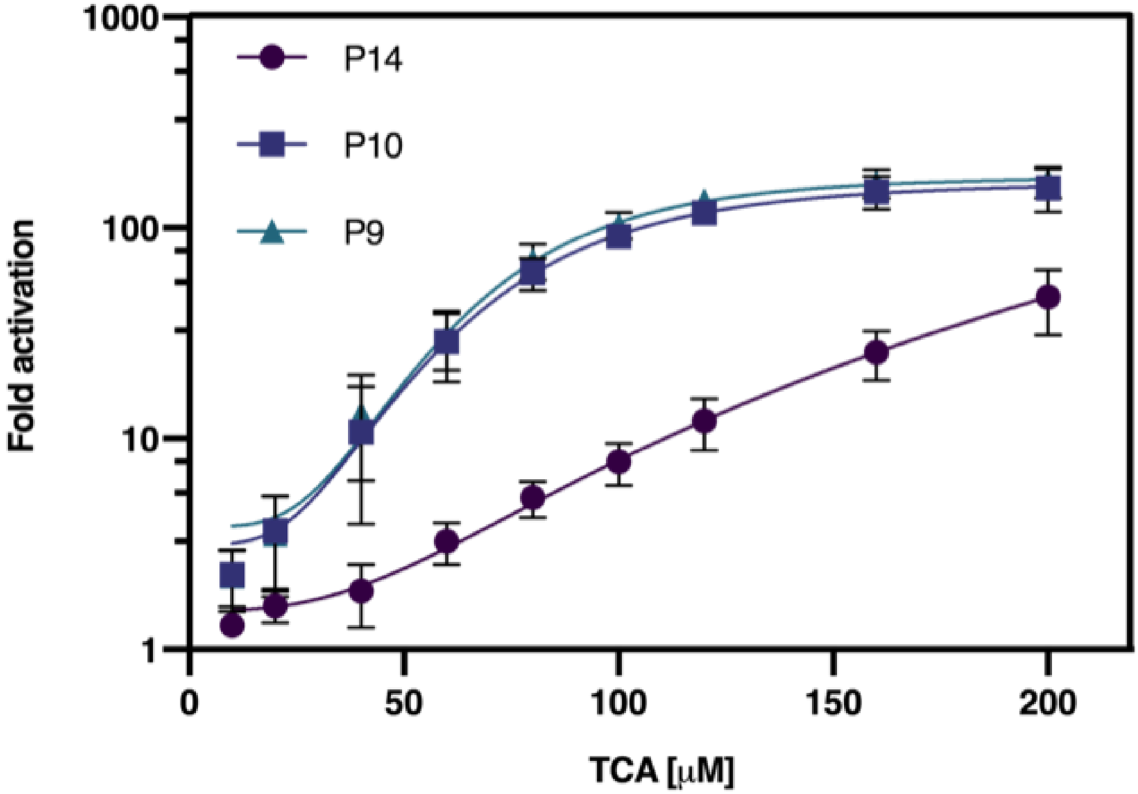

**Supplementary Figure 5: Activation fold of CadC-TcpP promoter variants (with P5-TcpH) responded to different concentrations of ligand TCA.**

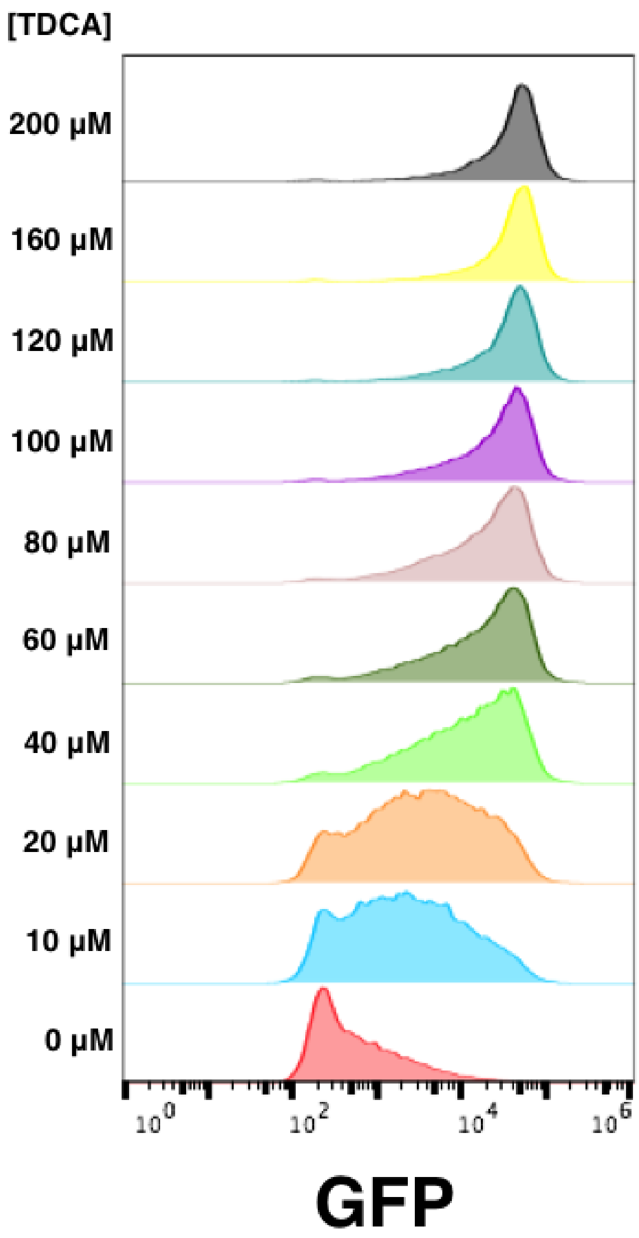

**Supplementary Figure 6: Response of P9-CadC-VtrA_P5-VtrC to different concentrations of TDCA.** Different concentration of TDCA used for titration are labeled at the left-y-axis.

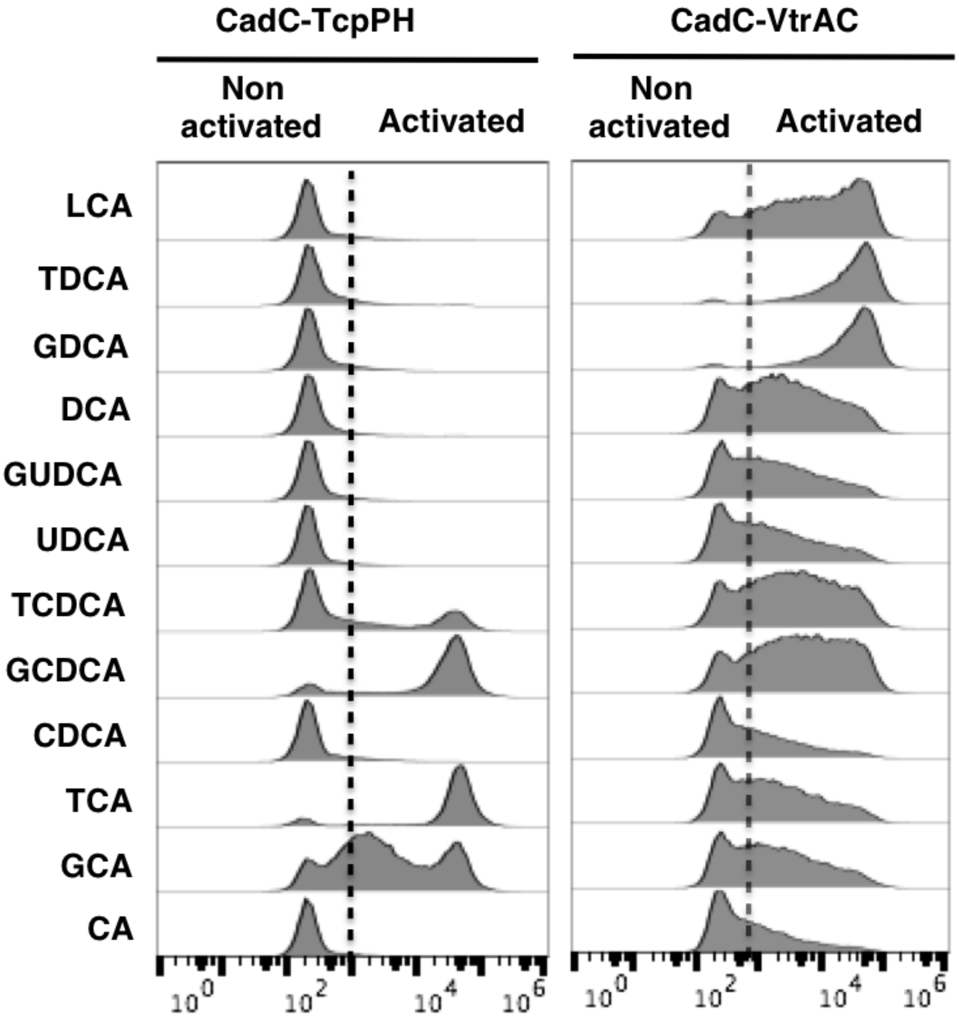

**Supplementary Figure 7: Response of CadC-TcpPH and CadC-VtrAC system to different type of bile salts.** The activated cells and non-activated cells are further distinguished by gating. The different bile salts used for profiling are labeled in the left-y-axis. Different from the TcpPH system which shows significant preference to primary conjugated bile salts, the VtrAC system shows different levels of preference to all bile salts. This result might reveal the potential of the hydrophobic inner chamber formed by VtrA/VtrC heterocomplex for the protein engineering of ligand specificity against different type of bile salts.

**a**

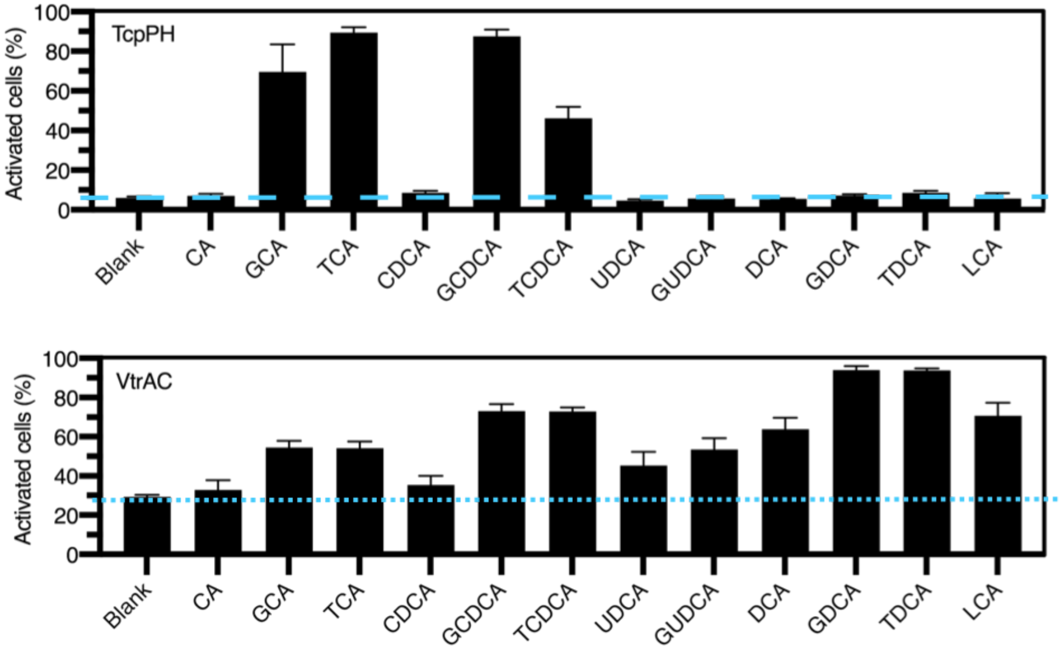

**b**

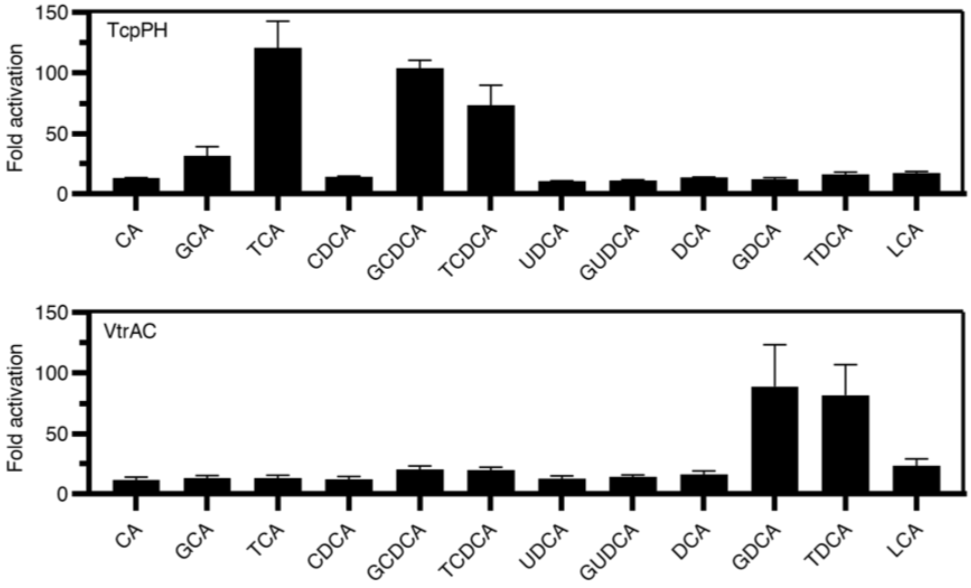

**Supplementary Figure 8: Response of CadC-TcpPH and CadC-VtrAC system to different bile salts are presented as (a) percentage of activated cells and (b) activation folds.** The population of activated cells are calculated through the gating shown in Fig. S8. The activation fold is calculated as the ratio of the geometric mean of activated cells divided by the geometric mean of non-activated cells.

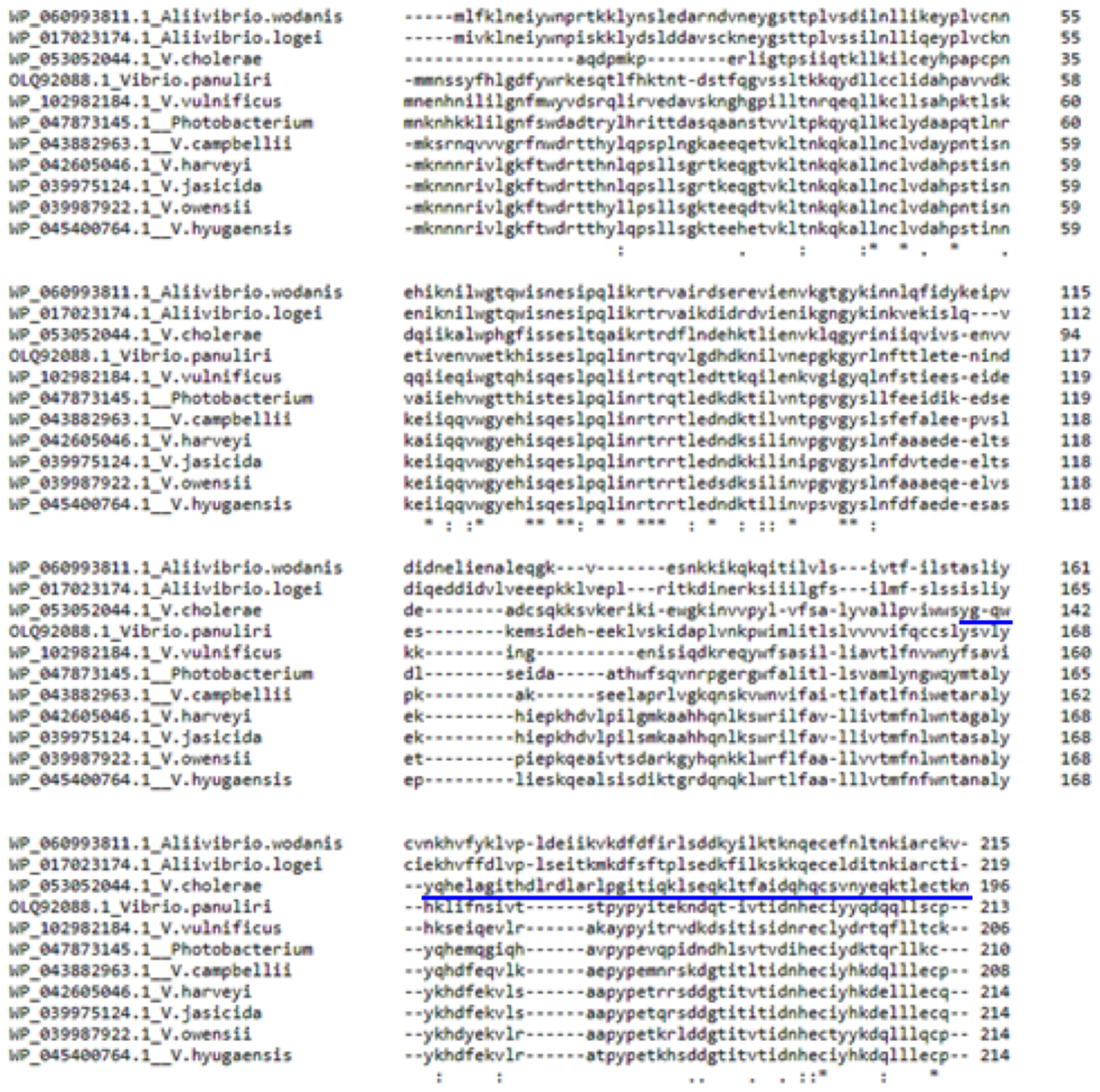

**Supplementary Figure 9: Multiple sequence alignment of the periplasmic domains from TcpP homologous proteins.** The periplasmic bile salt sensing domain of TcpP is highlighted by blue line.

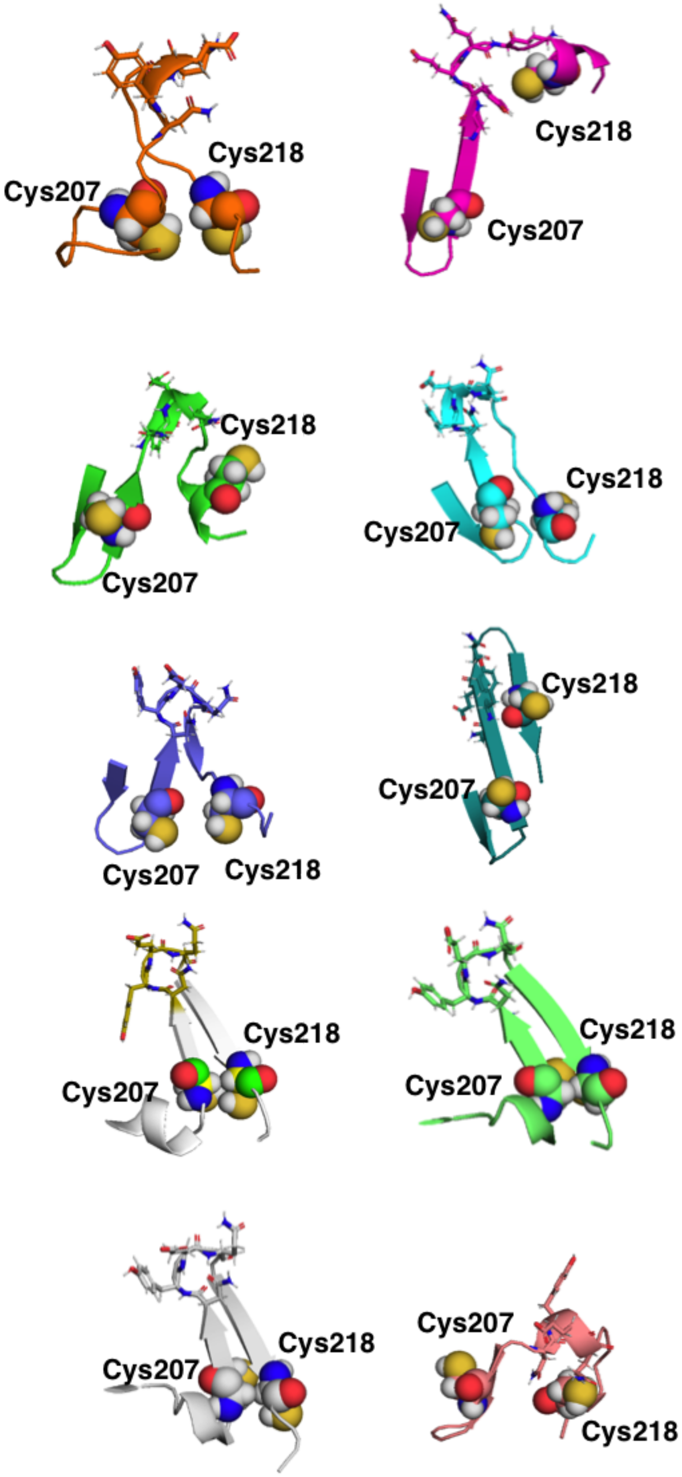

**Supplementary Figure 10: Molecular modeling of TcpP periplasmic sensing domain**

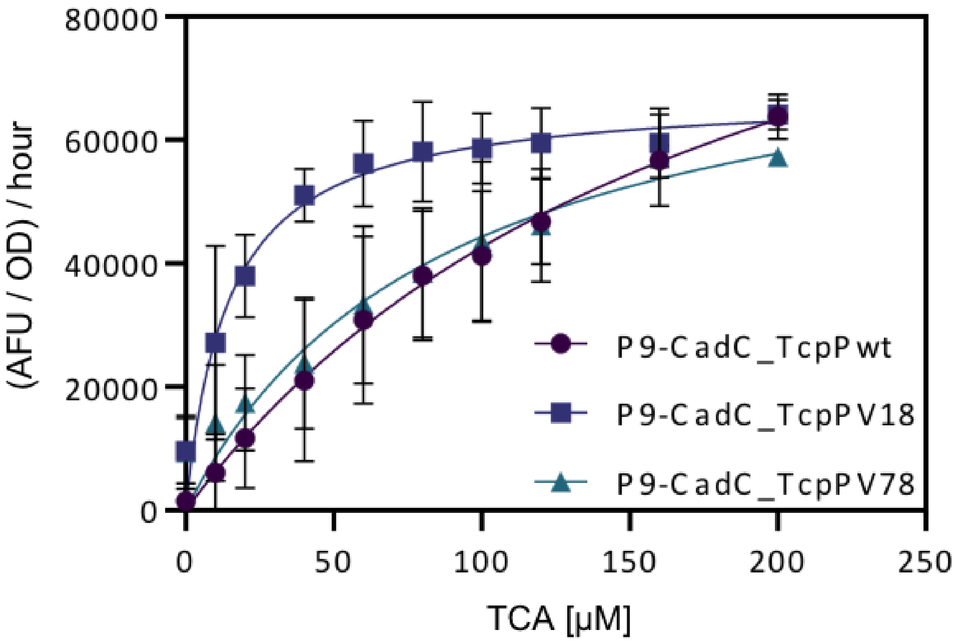

| | TcpPwt | TcpPV18 | TcpPV78 |
|---|---|---|---|
| **Vmax** | 123084 | 67529 | 83036 |
| **Km** | 188.8 | 14.44 | 87.4 |

**Supplementary Figure 11: Kinetics analysis of CadC-TcpPwt and CadC-TcpPmut-18**

**a**

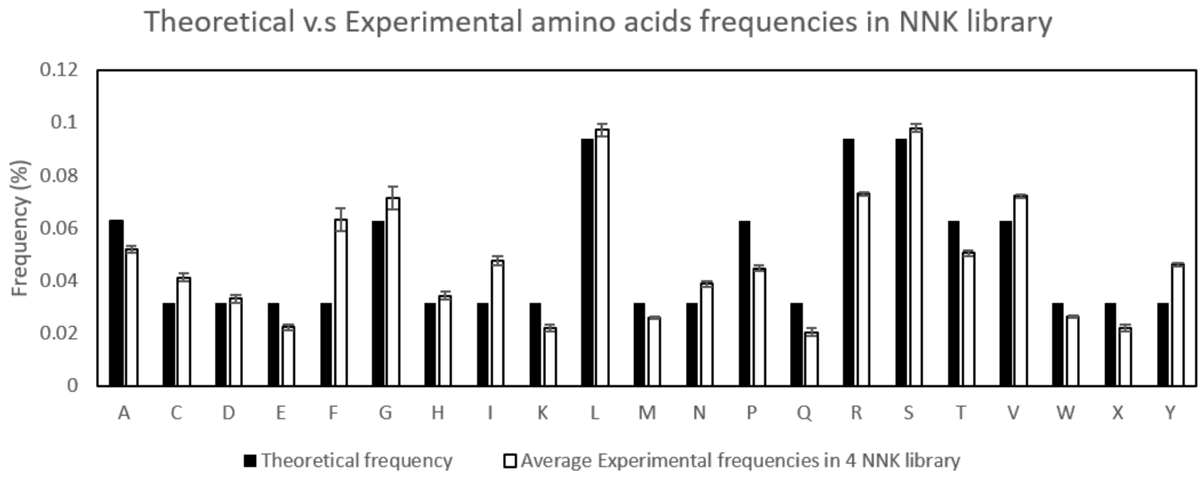

**b**

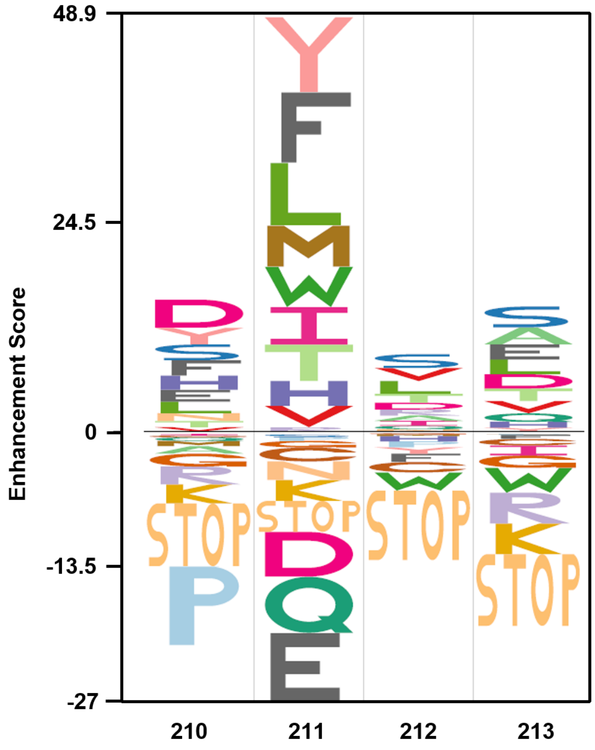

**Supplementary Figure 12: (a) Comparison of theoretical and experimental amino acid frequencies in preselected NNK library.** For verifying the quality of preselected TcpP synthesized 4 x NNK library, the summary counts of NGS results of each position were calculated as frequencies. The average values were further compared with theoretical frequencies of NNK library. **(b) The EDlogo plot of TcpP functional variants (from 3rd round of selection) compared with background frequencies (from preselected library) to highlight the enrichment and depletion score.** The y-axis of the logo is the enhancement score of each amino acid. STOP: the ‘TAG’ stop codon in NNK degenerate codons.

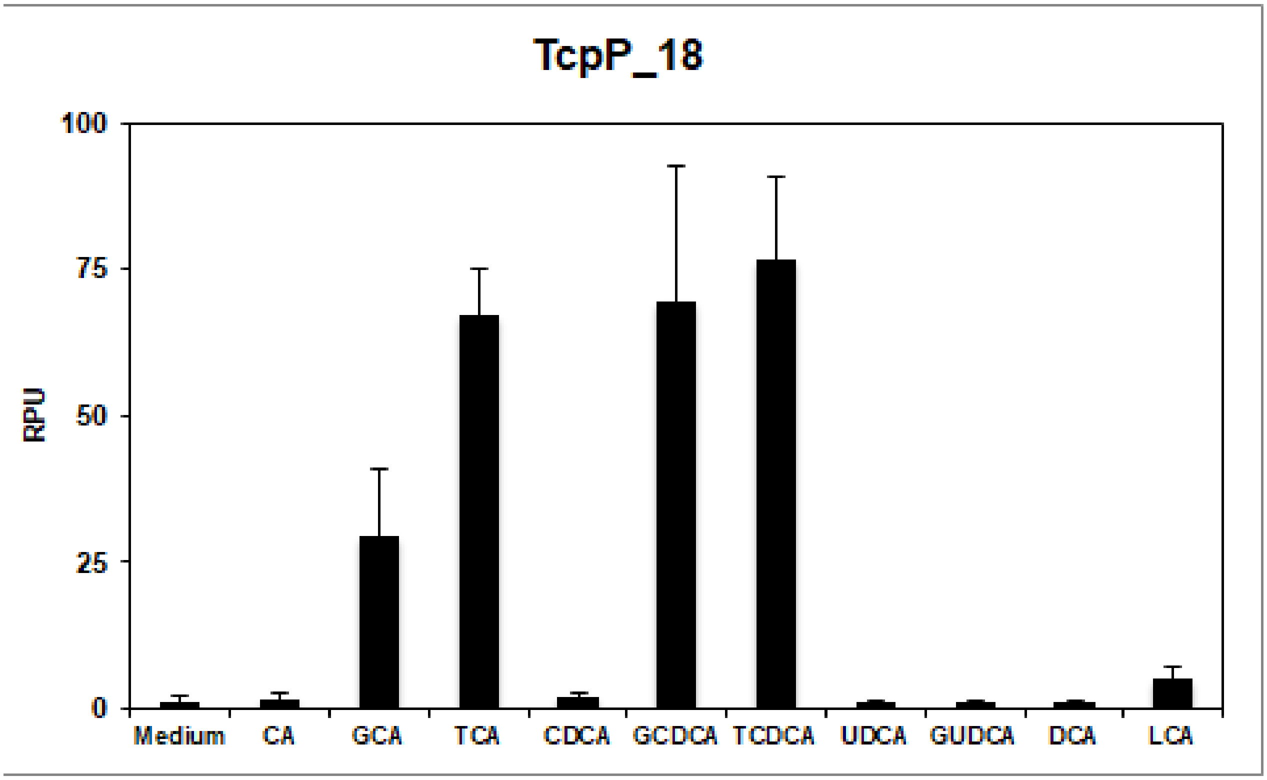

**Supplementary Figure 13: Bile salt response profile of TcpP18**

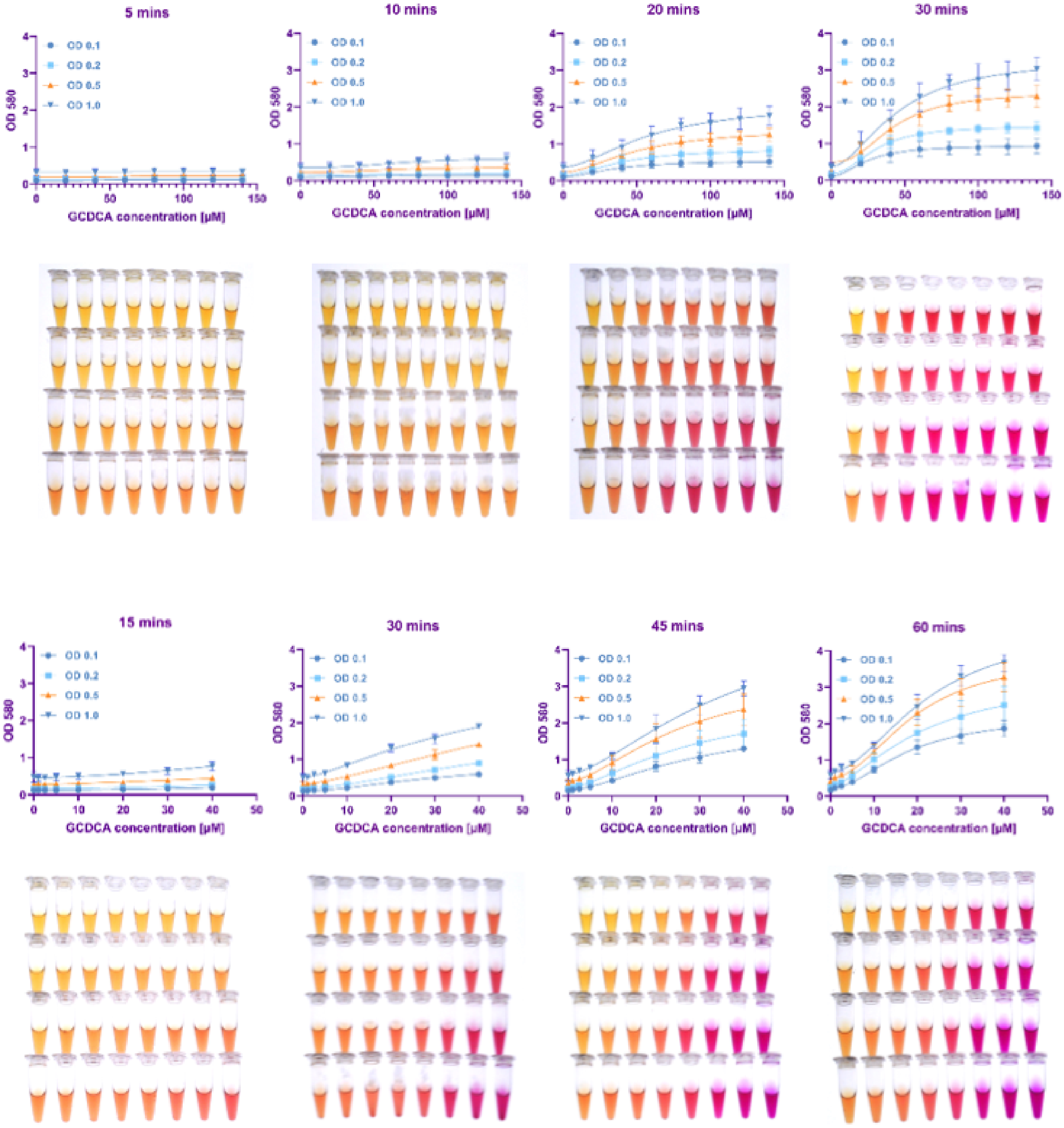

**Supplementary Figure 14: Fine tuning system performance of TcpP18-LacZ through varying cell number per reaction and sample incubation time.**

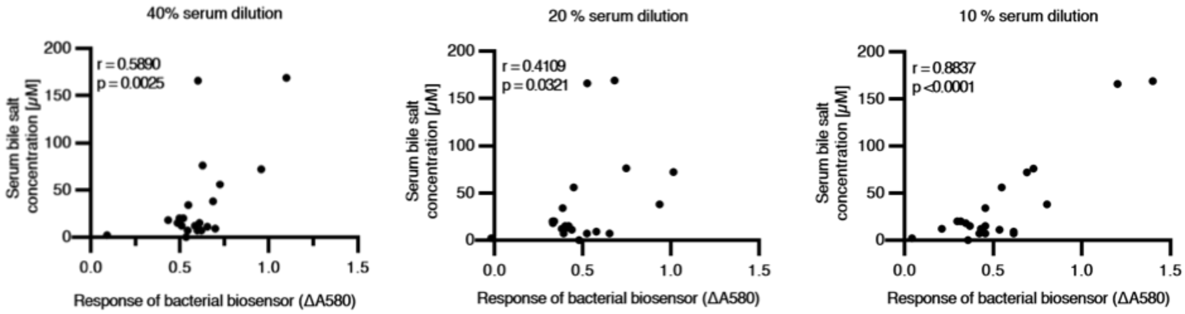

**Supplementary Figure 15: Determination of serum dilution rate in fresh bacterial biosensor cells.**

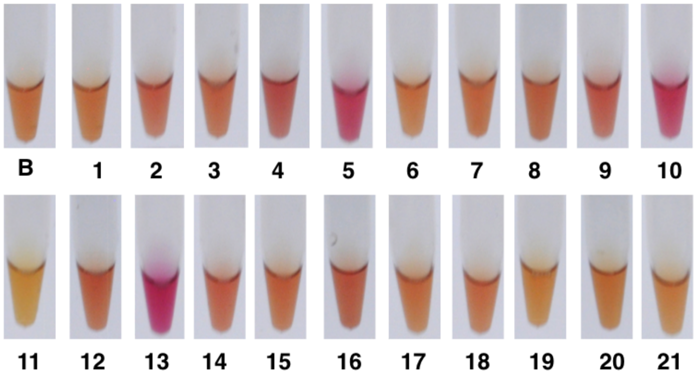

**# Blank = serum only**

| Sample | Blank | Patient 1 | Patient 2 | Patient 3 | Patient 4 | Patient 5 | Patient 6 | Patient 7 | Patient 8 | Patient 9 | Patient 10 |
|---|---|---|---|---|---|---|---|---|---|---|---|
| A580 | 0,77 | 0,80 | 0,89 | 0,86 | 0,98 | 1,52 | 0,83 | 1,01 | 0,82 | 1,11 | 1,23 |
| Stdeva | 0,05 | 0,12 | 0,06 | 0,06 | 0,14 | 0,17 | 0,10 | 0,19 | 0,13 | 0,17 | 0,10 |
| Sample | Patient 11 | Patient 12 | Patient 13 | Patient 14 | Patient 15 | Patient 16 | Patient 17 | Patient 18 | Patient 19 | Patient 20 | Patient 21 |
| A580 | 0,70 | 0,94 | 1,69 | 0,98 | 0,91 | 0,97 | 0,91 | 0,88 | 0,75 | 0,79 | 0,71 |
| Stdeva | 0,15 | 0,09 | 0,20 | 0,13 | 0,11 | 0,08 | 0,06 | 0,20 | 0,07 | 0,15 | 0,06 |

| Sample | | Patient 1 | Patient 2 | Patient 3 | Patient 4 | Patient 5 | Patient 6 | Patient 7 | Patient 8 | Patient 9 | Patient 10 |
|---|---|---|---|---|---|---|---|---|---|---|---|
| ΔA580 | (-Blank) | 0,04 | 0,13 | 0,10 | 0,22 | 0,75 | 0,06 | 0,25 | 0,05 | 0,35 | 0,47 |
| Sample | Patient 11 | Patient 12 | Patient 13 | Patient 14 | Patient 15 | Patient 16 | Patient 17 | Patient 18 | Patient 19 | Patient 20 | Patient 21 |
| ΔA580 | -0,07 | 0,17 | 0,92 | 0,22 | 0,15 | 0,20 | 0,15 | 0,11 | -0,02 | 0,02 | -0,06 |

**Supplementary Figure. 16 Visible color changes of bactosensor in clinical serum sample and measured absorbances values for each patient.**

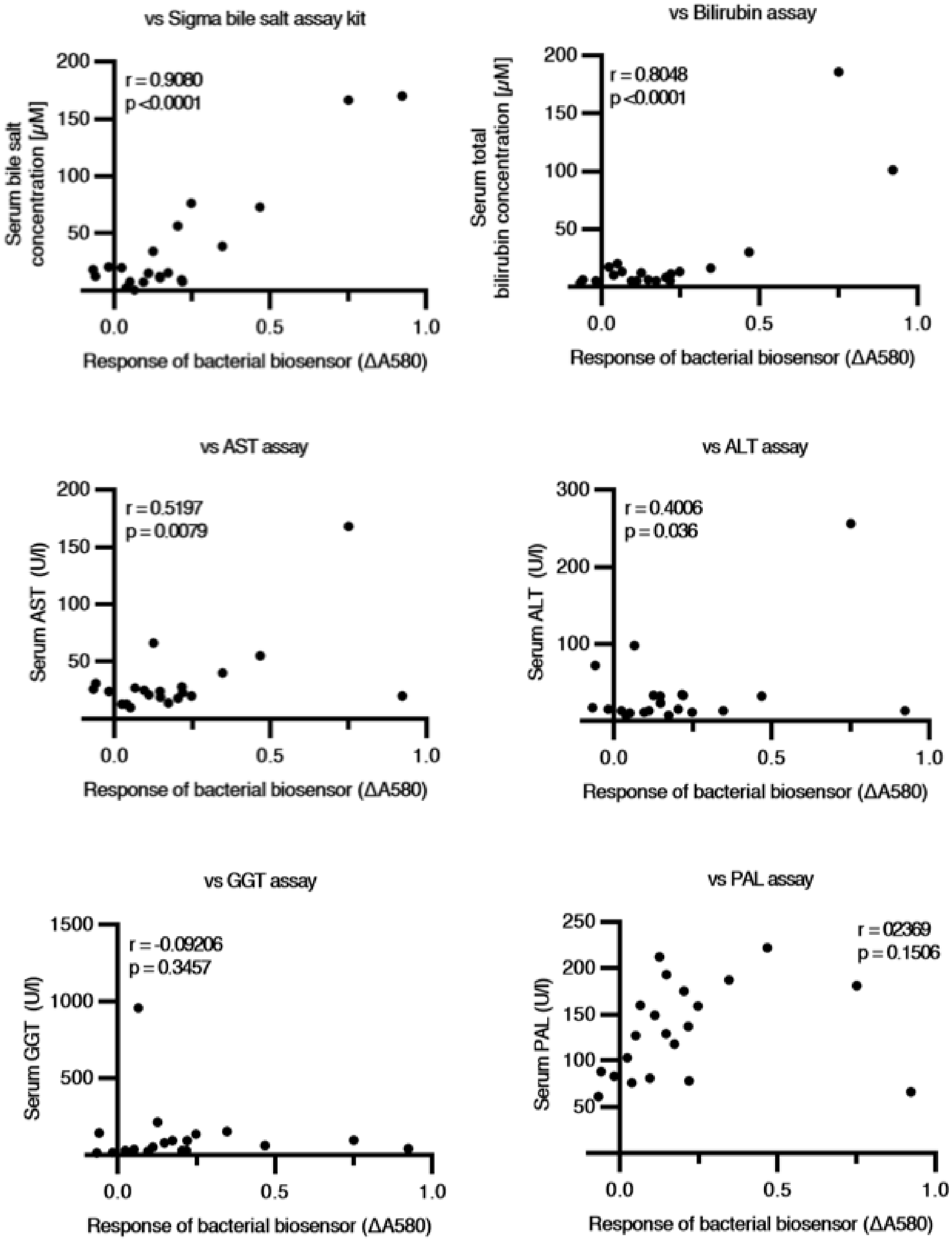

**Supplementary Figure. 17 Comparison of the data of bactosensor in clinical serum samples from 21 liver transplantation patients to different liver biomarkers.** These biomarkers includes bilirubin, aspartate aminotransferase (AST), alanine aminotransferase (ALT), gamma-glutamyl transferase (GGT), and alkaline phosphatase (ALP).

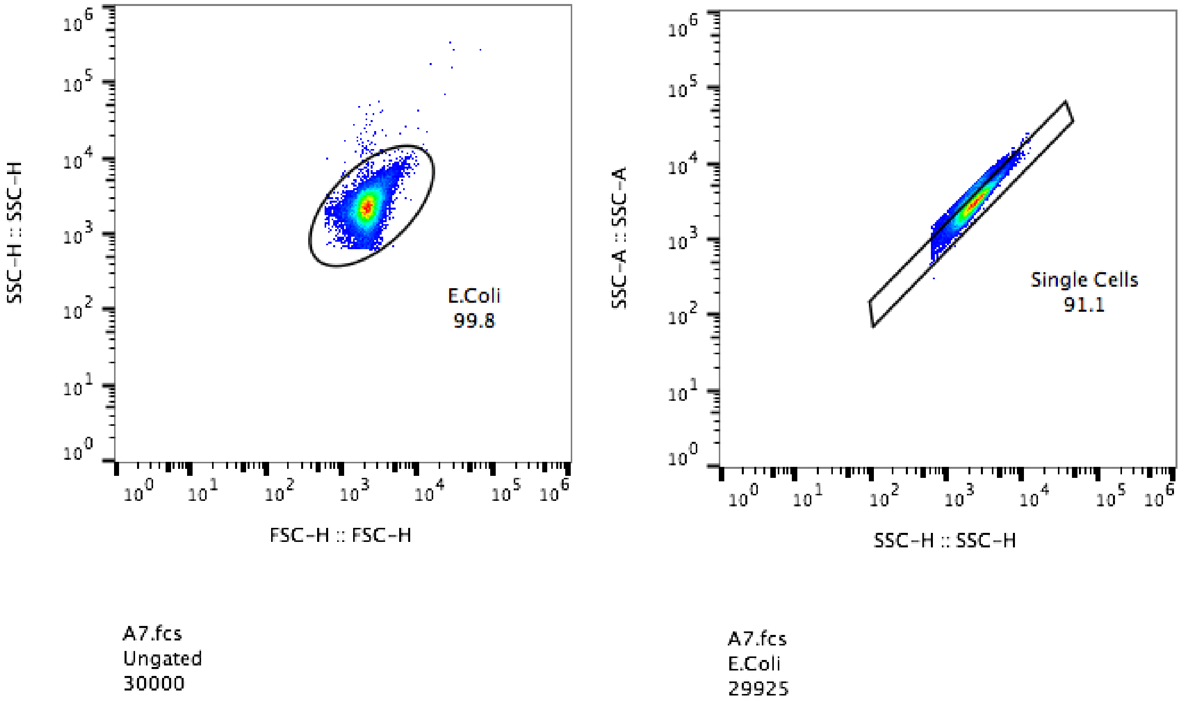

**Supplementary Figure 18: Figure exemplifying the flow cytometry gate strategy.** Gates were designed based on FSC-H vs SSC-H graphs to remove debris from the analysis (left pannel) and SSC-A vs SSC-H to doublet discrimination (right panel).

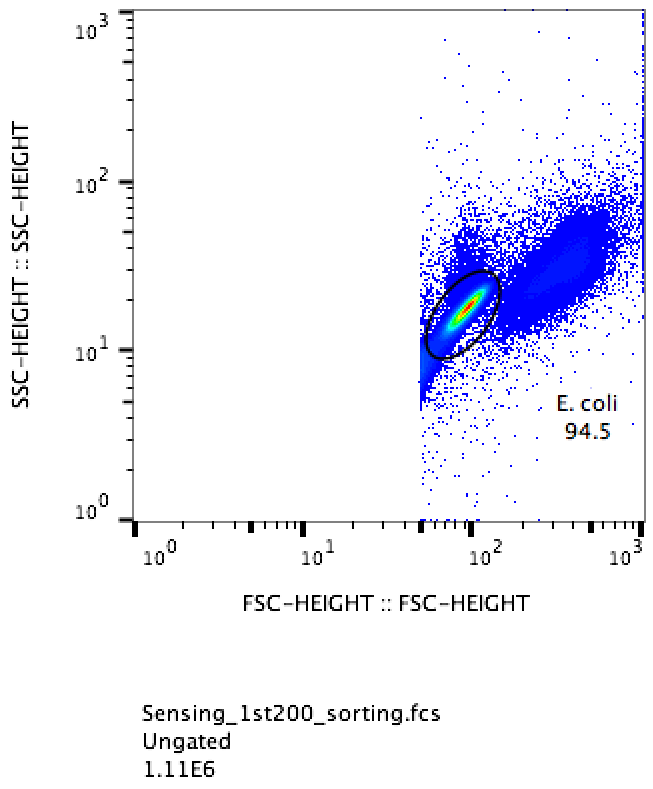

**Supplementary Figure 19: Figure exemplifying the cell sorter gate strategy.** Gates were designed based on FSC-H vs SSC-H graphs to remove debris from the analysis.

**Supplementary File1:** NGS summary counts of preselected synthetic NNK library and selected (Evo3) TcpP loop functional variants

**Table S1** Genetic parts

| Component | Nucleotide sequences |
|---|---|
| CadC DNA binding domain and juxtamembrane linker | ATGCAACAACCTGTAGTTCGCGTTGGCGAATGGCTTGTTACTCCGTCCATAAACCAAATTAGCCGCAATGGGCGTCAAC<br>TTACCCTTGAGCCGAGATTAATCGATCTTCTGGTTTTCTTTGCTCAACACAGTGGCGAAGTACTTAGCAGGGATGAACTT<br>ATCGATAATGTCTGGAAGAGAAGTATTGTCACCAATCACGTTGTGACGCAGAGTATCTCAGAACTACGTAAGTCATTAAAA<br>GATAATGATGAAGATAGTCCTGTCTATATCGCTACTGTACCAAAGCGCGGCTATAAATTAATGGTGCCGGTTATCTGGTACA<br>GCGAAGAAGAGGGAGAGGAAATAATGCTATCTTCGCCTCCCCCTATACCAGAGGCGGTTCCTGCCACAGATTCTCCCT<br>CCCACAGTCTTAACATTCAAAACACCGCAACGCCACCTGAACAATCCCCAGTTAAAAGCAAACGA |
| TcpP transmembrane domain and periplasmic domain | GTGGTGCCGTATCTGGTGTTTTCAGCGTTATATGTTGCCTTATTACCTGTCATTTGGTGGTCATACGGGCAATGGTATCAA<br>CACGAATTGGCCGGGATTACTCACGATTTACGTGACTTAGCCCGTCTTCCTGGAATCACAATCCAGAAACTTAGCGAGC<br>AGAAGTTGACCTTTGCTATCGACCAGCATCAATGTTCGGTGAATTACGAACAAAAGACGTTGGAATGTACTAAGAACTAAT<br>AA |
| TcpH | ATGCACAAGAAGTTAAAGGCTTGGGGAGGGGCCACTGGTCTGTTTGTGGTCGCCCTGGGGGTAACAATTATTGCTCTT<br>CCTATGCGCCAGAAAAATAGCCACGGGACCATGATTATTGATGGAACCGTTACTCAAATCTTTAGCACCTACCAGGGCAA<br>TCTGTCTAATGTGTGGCTTACTCAGACGGACCCTCAAGGTAATGTTGTTAAATCATGGACTACACGCTACCAAACATTGC<br>CTGACCCTTCATCACAAAAGCTGAATCTGATCCCCGATTACTCCCAATCGAATGCTTCTCGTGATTACAATGTATTGTCGA<br>TCTATCAACTTGGCAAAGGGTGCTTTCTGGCTTTTCCTTACAAATTGCTGACGGCGGAAAAGATGTGGTTCTCGTGTCA<br>GAGCGATTTCTAA |
| VtrA transmembrane domain and periplasmic domain | ATTCTTCTTATCAGCATTATTTTGCAGTTGGCTTTCATCATCTATGTGGCTTATAGCTACACCAGTATTTTCGTGTCGTCCAC<br>CGCAAAGGACGACTATCCCAGTTTATCGTTCCAACAAGACTACGTCTACATCTTTTCTTCAGACTTTCAATTAAGCGAGG<br>AGCTGGGCGTGGCCCTTATCAACGCTTTAAGTGCAAAAGAAATTGTACCCGAGCGTCTTTACGTTATGTTGAATGACAAG<br>ACCATCTCGTTCAGTTTTATCAGCAAGAATAAGAAGTCCAAGAATCGCGTATTAAGCACGGAAAAGAAACTCAATTATAAG<br>CATATCTCCGAGTACATCGTCAATGAGATTGAGTACTAATAA |
| VtrC | ATGAAACTTAATATTAAGCGTCTGCATTTGAGCCTGACCCTGATGAGTGTCGTCATGCTGCTGGTAATCATCTACAATAATT<br>TCTTTCAACCCGTGCACTTCTACGAAACGTCATACAAGTACCAAGCGGCGGACTCGACGTACATGCACGACGTCGCGA<br>TCAATGTCTCGATTAAGGGCAACCACTTCACCTCAGATATCATCATCCGCGAGCTGGTGAAGTCAGAGAACAAGAATTAC<br>TATAACGTTATTGGACATGGAGACATCATTCAAAAGAACACCCATCAATACTACTTGAACTTCGATAACATCGACGTTTACA<br>CGGGTACTAATAAAGCGAACATGAAGCCGTATAAGGAACCGACTAGCATCTCCTCGCTCATCAACAAGTCAAATAACATT<br>CGCGTTGTTTACTTATCGGAAGAGTATGTTGTGGTAGAGTTCTTCTTTTATGATGGACAGATTATCACATTGCATCGCTATTA<br>ATAA |
| sfGFP | ATGTCAAAAGGAGAAGAACTTTTTACAGGTGTAGTACCTATCTTGGTTGAATTGGATGGTGATGTTAACGGTCACAAATTT<br>TCTGTACGTGGTGAAGGTGAAGGTGATGCAACTAACGGTAAATTGACACTTAAATTCATTTGTACAACTGGAAAACTTCCT<br>GTTCCTTGGCCTACTCTTGTTACAACATTGACATATGGAGTACAATGTTTTTCACGTTATCCTGATCATATGAAACGTCACG<br>ATTTTTTTAAATCTGCTATGCCAGAAGGTTATGTACAAGAACGTACAATTTCATTTAAAGATGACGGAACATATAAAACACGT<br>GCTGAAGTAAAATTCGAAGGTGACACTCTTGTTAATCGTATCGAATTGAAAGGAATCGATTTCAAAGAAGATGGTAACATT<br>TTGGGACACAAACTTGAATACAACTTCAACTCTCATAATGTTTATATCACAGCTGACAAACAAAAAAACGGTATTAAAGCTA<br>ATTTTAAAATTCGTCACAATGTTGAAGATGGATCTGTTCAATTGGCTGATCATTATCAACAAAATACACCAATCGGAGACGG<br>ACCAGTATTGCTTCCAGATAACCACTACCTTTCTACTCAATCAGTTCTTTCAAAAGATCCTAACGAAAAACGTGACCATAT<br>GGTACTTCTTGAATTTGTTACAGCAGCAGGTATCACTCACGGTATGGACGAACTTTATAAATAA |
| **Promoters** | **Nucleotide sequences** |
| P14 | TTGACAATTAATCATCCGGCTCGTATAATGTGTGGA |
| P10 | TTTCAATTTAATCATCCGGCTCGTATAATGTGTGGA |
| P9 | TTGCCTCTTAATCATCGGCTCGTATAATGTGTGGA |
| pCadBA | ATCCATTGTAAACATTAAATGTTTATCTTTTCATGATATCAACTTGCGATCCTGATGTGTTAATAAAAAACCTCAAGTTCTCAC<br>TTACAGAAACTTTTGTGTTATTTCACCTAATCTTTAGGATTAATCCTTTTTTCGTGAGTAATCTTATCGCCAGTTTGG |

**Table S2-5 Patient characteristics**

**Table S2.** Patients characteristics (n=21)

| **Characteristics** | **Value** |
|---|---|
| **Age (years)** | 57.5±10.4 |
| **Male, n (%)** | 15 (71.4) |
| **Underlying liver disease** | |
| Alcoholic cirrhosis, n (%) | 7(33.3) |
| Hepatocellular carcinoma, n (%) | 4 (19) |
| Virus related cirrhosis (HBV, HCV), n (%) | 2 (9.5) |
| Primary sclerosing cholangitis or primitive biliary cholangitis | 5 (24) |
| Other (non-alcoholic steato-hepatitis, Budd-Chiari,auto immune hepatitis) | 3 (14.2) |
| **Time from liver transplantation to serum analysis in months, median (SD)** | 8± 38 |
| **Still alive at the end of the study ,** n (%) | 18 (85.7) |

**Table S3.** Patients characteristics (n=21)

| Patient number | Age range | Sex | Cause of transplant | Year of transplant |
|---|---|---|---|---|
| 1 | 50-69 | m | Alcoholic | 2020 |
| 2 | 60-69 | f | NASH | 2019 |
| 3 | 60-69 | m | HCC | 2019 |
| 4 | 60-69 | m | Alcoholic | 2019 |
| 5 | 30-39 | m | vascular cirrhosis | 2020 |
| 6 | 60-69 | m | HCV | 2006 |
| 7 | 40-49 | f | auto-immune acute hepatitis | 2020 |
| 8 | 60-69 | m | PSC | 2020 |
| 9 | 60-69 | f | cirrhosis NASH | 2020 |
| 10 | 40-49 | f | PSC | 2014 |
| 11 | 60-69 | f | autoimmune cirrhosis | 2019 |
| 12 | >70 | m | budd chiari disorder | 2019 |
| 13 | 60-69 | m | alcoholic cirrhosis | 2013 |
| 14 | 30-39 | m | PSC | 2019 |
| 15 | >70 | m | NASH | 2019 |
| 16 | 60-69 | m | ischemic cholangitis | 2019 |
| 17 | 50-59 | m | HVC/HCC/alcoholic cirrhosis | 2019 |
| 18 | 50-59 | f | HVC/HCC/alcoholic cirrhosis | 2019 |
| 19 | 50-59 | m | HBV | 2018 |
| 20 | 60-69 | f | PSC | 2020 |
| 21 | 40-49 | f | HBV/HCV/HDV | 2016 |

PSC: Primary Sclerosing Cholangitis
HBV: Hepatitis B Virus
HCV: Hepatitis C Virus
HCC: Hepato Cellular Carcinoma
NASH: Non-Alcoholic SteatoHepatitis

**Table S4.** Patients characteristics (n=21)-Hepatic tests and bactosensor measurements

| Patient number | ASAT | ALAT | GGT | Pal | Total bilirubin | hospital bile salt | Bile salt (sigma kit) | BacSensor (OD 580) |
|---|---|---|---|---|---|---|---|---|
| 1 | 13 | 7 | 18 | 76 | 10 | 4,7 | 2,56 | 0,07 |
| 2 | 66 | 33 | 214 | 212 | 12 | 24 | 34,44 | 0,16 |
| 3 | 25 | 11 | 24 | 81 | 5 | 7,5 | 7,59 | 0,13 |
| 4 | 23 | 33 | 94 | 78 | 11 | 5,4 | 7,86 | 0,32 |
| 5 | 168 | 256 | 96 | 181 | 186 | 193,1 | 166,21 | 0,91 |
| 6 | 27 | 98 | 959 | 160 | 13 | 2,7 | 0,62 | 0,06 |
| 7 | 20 | 11 | 136 | 159 | 13 | 19 | 76,22 | 0,43 |
| 8 | 10 | 10 | 36 | 127 | 20 | 7,9 | 7,86 | 0,16 |
| 9 | 40 | 13 | 154 | 187 | 16 | 25 | 38,74 | 0,51 |
| 10 | 55 | 32 | 60 | 222 | 30 | 55,8 | 72,91 | 0,40 |
| 11 | 26 | 17 | 14 | 61 | 3 | 10,4 | 18,24 | 0,05 |
| 12 | 14 | 7 | 94 | 118 | 5 | 13,2 | 15,65 | 0,16 |
| 13 | 20 | 13 | 41 | 66 | 101 | 77 | 169,91 | 1,11 |
| 14 | 28 | 34 | 28 | 137 | 5 | 10,6 | 9,63 | 0,32 |
| 15 | 19 | 23 | 78 | 193 | 6 | 11,3 | 11,10 | 0,24 |
| 16 | 18 | 15 | 29 | 175 | 8 | 40,9 | 56,58 | 0,25 |
| 17 | 24 | 32 | 81 | 129 | 5 | 8,4 | 12,07 | 0,14 |
| 18 | 21 | 13 | 50 | 149 | 5 | 11,2 | 15,37 | 0,07 |
| 19 | 24 | 15 | 17 | 83 | 5 | 3,6 | 20,71 | 0,00 |
| 20 | 13 | 13 | 28 | 103 | 17 | 12,6 | 20,08 | 0,02 |
| 21 | 31 | 72 | 143 | 88 | 6 | 3,2 | 12,60 | -0,08 |

**Table S5.** Patients acute clinical conditions

| Patient number | Acute clinical condition |
|---|---|
| 1 | follow up |
| 2 | bypass/switch duodéno pancréas |
| 3 | follow up |
| 4 | type 2 diabetes |
| 5 | transplant liver and kidney, graft rejection/ biliary stenosis |
| 6 | graft rejection and biliary duct obstruction/stenting 48 H before blood test |
| 7 | Kehr drain |
| 8 | follow up |
| 9 | sepsis (acute general infection) and hepatic artery stenosis |
| 10 | follow up and new cirrhosis |
| 11 | fiever |
| 12 | ischemic cholangitis |
| 13 | type 2 diabetes |
| 14 | follow up |
| 15 | follow up |
| 16 | follow up, phosphatase alcaline d'origine osseuse, portage BLSE urinaire |
| 17 | follow up and bile duct stenosis |
| 18 | follow up |
| 19 | acute colic infection |
| 20 | follow up |
| 21 | angiocholitis |
